# The complete diploid reference genome of RPE-1 identifies human phased epigenetic landscapes

**DOI:** 10.1101/2023.11.01.565049

**Authors:** Emilia Volpe, Luca Corda, Elena Di Tommaso, Franca Pelliccia, Riccardo Ottalevi, Danilo Licastro, Andrea Guarracino, Mattia Capulli, Giulio Formenti, Evelyne Tassone, Simona Giunta

**Author notes:** Correspondence: Simona Giunta.

## Abstract

Comparative analysis of recent human genome assemblies highlights profound sequence divergence that peaks within polymorphic loci such as centromeres. This raises the question about the adequacy of relying on human reference genomes to accurately analyze sequencing data derived from experimental cell lines. Here, we generated the complete diploid genome assembly for the human retinal epithelial cells (RPE-1), a widely used non-cancer laboratory cell line with a stable karyotype, to use as matched reference for multi-omics sequencing data analysis. Our RPE1v1.0 assembly presents completely phased haplotypes and chromosome-level scaffolds that span centromeres with ultra-high base accuracy (>QV60). We mapped the haplotype-specific genomic variation specific to this cell line including t(X*q*;10*q*), a stable 73.18 Mb duplication of chromosome 10 translocated onto the microdeleted chromosome X telomere t(X*q*;10*q*). Polymorphisms between haplotypes of the same genome reveals genetic and epigenetic variation for all chromosomes, especially at centromeres. The RPE-1 assembly as matched reference genome improves mapping quality of multi-omics reads originating from RPE-1 cells with drastic reduction in alignments mismatches compared to using the most complete human reference to date (CHM13). Leveraging the accuracy achieved using a matched reference, we were able to identify the kinetochore sites at base pair resolution and show unprecedented variation between haplotypes. This work showcases the use of matched reference genomes for multi-omics analyses and serves as the foundation for a call to comprehensively assemble experimentally relevant cell lines for widespread application.

**Highlights:**

- We generated the complete phased genome assembly of one of the most widely used non-cancer cell lines (RPE-1) with a stable diploid karyotype
- We used this genome as a matched reference to analyze sequencing data from RPE-1
- Mapping to the RPE1v1.0 genome improves alignment quality, faithful assignment of reads to each haplotype, and epigenome peak calling accuracy uncovering inter-haplotype variation
- Use of the matched reference genome enables epigenetic precision in identifying for the first time the kinetochore site at base pair resolution for each haplotype
- The RPE-1 genome represents a new telomere-to-telomere (T2T) human diploid reference for the scientific community that will advance genetic and epigenetic research across fields using this cell line

## INTRODUCTION

Recent advances in DNA sequencing and genome assembly have led to the achievement of complete human genomes. These include the near-homozygous CHM13 ^1^ and genomes from individuals of diverse ancestry ^2,3^. These and others that have since been added ^4–6^ represent an important step toward a more equitable representation of human genomic diversity ^3,7^. Comparative analyses of the currently available assemblies have revealed an unexpected level of sequence variation ^3,8^, including inter-individuals divergence ^3^ and population-level variants ^9–11^. Differences within an individual genome pertaining to the parental alleles are largely unexplored ^2,12^ due to lack of high-quality human diploid genomes with fully phased haplotypes. Early evidence of the maternal and paternal haplotypes in HG002 show remarkable sequence divergence was shown between, with non-synonymous amino acid changes in almost 48% protein-coding genes that peaks at polymorphic loci ^2^, in contrast with previous evidence pointing to low haplotype variation ^13^. This profound genomic diversity ^3^ underlies phenotypic traits ^14–16^ and significantly affects targeted gene editing and in turn, clinical outcomes ^17^. The extent of sequence variation highlights the need to reconsider our current approach based on reference genomes. In particular, the analysis of multi-omics sequencing data isolated from laboratory cell lines presents challenges when using a non-matched reference, the so-called ‘reference bias’ ^18^. These challenges are particularly important for faithful studies of our most polymorphic and divergent regions including centromeres ^8^. Human centromeres consist of repetitive monomers of alpha-satellite DNA hierarchically organized into near-identical reiterating higher order repeats (HORs) arrays ^19–22^. While centromere DNA varies between individuals, their essential function in chromosome segregation is epigenetically supported by the centromere-specific histone H3 variant CENP-A ^23,24^. Two features point to the kinetochore binding site within the active HORs: a sudden drop in DNA methylation named centromere dip region (CDR) ^22^, and increased density of CENP-A nucleosomes estimated to reach ∼1 every 4 canonical nucleosomes compared to 1 every 20 canonical nucleosomes in the rest of the active region ^25^. In spite of advances in computational tools and long-reads sequencing technologies ^1,22,26–29^, reliable assembly of centromeres for all chromosomes remains a challenge ^30^. In fact, centromeres were largely omitted in the recent human pangenome draft ^3^, with no complete diploid human reference genome of experimentally amenable cell lines available to date. Recent genetic and epigenetic characterization of complete centromeres leveraged near-homozygous *ad hoc* cell lines derived from anucleate fertilized oocyte of non-viable molar pregnancy ^1,22,31^, that do not allow for the assessment of inter-haplotype polymorphism within the same diploid individual. While not experimentally amenable, these immortalized cell lines lack complications from allelic variation facilitating complete assembly ^26,31–33^ and centromere comparison ^21,34^. Here, we present the first to our knowledge high-quality human diploid reference genome from human cell line amenable to laboratory experimentation that can be used to interrogate genetic and epigenetic changes between genomes and haplotypes. The retinal pigment epithelial line (RPE-1) is one of the most commonly used non-cancer laboratory cell lines, counting thousands of scientific publications ^35^. Because of their stable diploid karyotype and extensive sequencing data publicly available, we chose this cell line to generate a new telomere-to-telomere (T2T) human diploid genome assembly with phased haplotypes and used it to baseline variation at centromeres. Using this assembly as diploid reference genome drastically improves alignments of RPE-1 reads from multi-omics experiments (Fig. 1a). We describe this novel approach to match reads with the assembly as ‘*isogenomic*’ reference genome. Our work represents a proof-of-concept that calls for a comprehensive catalog of complete genome assemblies for commonly used cells, including diploid embryonic and induced pluripotent stem cells (ESC and iPSC), primary and disease lines, for a widespread application of *isogenomic* reference genomes to enable faithful multi-omics analyses and high-precision epigenetics.

**Figure 1.**
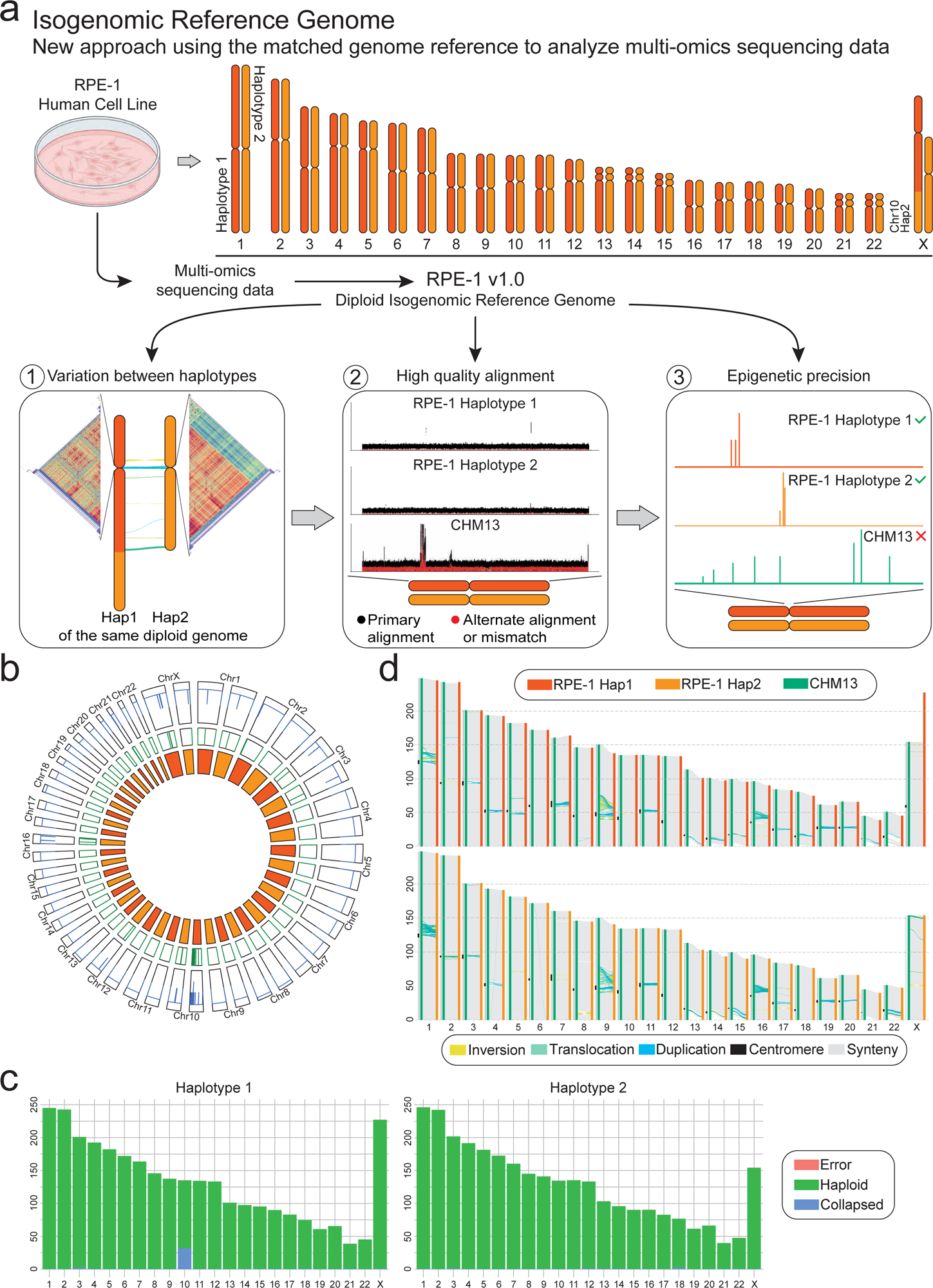
The diploid reference genome of RPE-1 enables faithful multi-omics data analysis. (a) Schematic of the workflow from *in vitro* experiments to multi-omics data analyzed using the isogenomic reference genome (top). The advantages of this model are 1) the ability to assess the variation between haplotypes, 2) improved whole-genome alignment, and 3) faithful phased epigenomic peaks calling (bottom). (b) Regional and structural assembly errors identification using alignment Clipping information. Circos plot shows Assembly Quality Indicators (AQI), Clip-based Regional Errors (CREs) at precise location, for each chromosome of both haplotypes (from outside to inside). (c) Synteny between all chromosomes of RPE-1 haplotypes (Haplotype 1, Hap 1 and Haplotype 2, Hap 2) and CHM13 (haploid). RPE-1 shows comparable chromosome length to CHM13, with the majority of the variation in centromeric and pericentromeric regions. (d) Reliability of the RPE-1 chromosomes using read mapping. Regions flagged as ‘haploid’ are error-free (green).

## RESULTS

### A telomere-to-telomere human diploid reference genome for the RPE-1 cell line

To improve the standard practice of aligning sequencing reads originated from laboratory cell lines against a human reference genome from a different individual, we present a new approach using the matched or ‘*isogenomic*’ reference genome (Fig. 1a). This approach enables (1) to assess differences between haplotypes, (2) optimal alignment and (3) for the first time, phased epigenetic mapping at base pair resolution. To achieve this, we generated the first complete *de novo* diploid assembly of one of the most commonly used non-cancer laboratory cell lines for multi-omics experiments, the human retinal pigment epithelial cells RPE-1 (Fig. 1a). We produced state-of-the-art sequencing data (Table 1, Supplementary Fig. 1-2, Methods). After comparing two *de novo* whole-genome assemblers, Verkko ^12,36^ (Supplementary Fig. 3a-b) and Hifiasm ^37^ (Supplementary Fig. 3c), we generated the final assembly using Verkko (Supplementary Note 1). To achieve complete phasing of the RPE-1 diploid genome in absence of parental information for trio-binning ^2^, we produced Hi-C data integrated onto the unphased Verkko’s output to obtain haplotype 1 (Hap 1) and haplotype 2 (Hap 2) and to guide resolution of possible missed joins by manual dual curation using the Hi-C contact map ^2^ (Supplementary Fig. 4a-c, Supplementary Note 2, Methods). A variety of evaluation tools ^38–40^ confirmed the high quality of the final RPE-1 assembly (i.e. HPRC, T2T, HGSVC), with estimated base accuracy over 99.9999%, specifically a QV of 64.1 on Hap 1 and 61.8 on Hap 2, (Table 1) and 99.8% completeness without errors for both haplotypes ^3,41^ (Fig. 1b). Particularly, 6.01 Gb corresponds to the reliable genome flagged as haploid, 2.19 Mb were errors and 53 Mb collapsed regions (Fig. 1c). Each haplotype of RPE-1 human genome, as expected, was syntenic whole-genome compared to CHM13 (Fig. 1d), coherent with the absence of structural assembly errors.

The RPE-1 total genomic content was 3.06 Gb and 2.99 Gb for Hap 1 and Hap 2, and chromosomes length was comparable to the previous genome assemblies CHM13, HG38 (GRCh38.p14) and HG002 (HG002v0.7). The only exception is for a copy of RPE-1 chromosome X of Hap 1 with a total length of 227.21 Mb that deviates from the expected 155 Mb (Supplementary Fig. 5a-b). Notably, this is not a *de novo* somatic rearrangement but corresponds to a stable marker chromosome for the RPE-1 cell line that we ^42,43^ and others ^35,44–46^ had previously observed by cytological and sequencing analyses. The chromosome contacts between X and 10 were also flagged on the dual Hi-C contact map for both haplotypes (Supplementary Fig. 3a), leading an increase in gaps for chromosome 10 during assembly step, showing an increase in coverage in the string graph nodes confirmed a duplicated 10*q* (Supplementary Fig. 5c-d). Also, the interlinked triplex bandage of chromosome X *q*-arm fused to chromosome 10 *q-*arm is present on the DeBrujin graph of the unphased assembly, but not upon downsampling of Verkko with HiFi and ONT-UL over 100 kb alone (Supplementary Fig. 3b), striking evidence of a major long-range chromosomal translocation automatically detected by a genome assembler on a diploid human genome (Supplementary Fig. 5e).

Ongoing chromosomal rearrangements in laboratory cell lines can represent a challenge in defining a consensus reference. To confirm that RPE-1 are a karyotypically stable diploid cell line, we cytogenetically analyzed variations across the cell population, in different batches and during passages in culture As expected, we did not observe overt karyotypic variations at the cytogenetic level, including stable presence of the t(Xq;10q) marker chromosome in all batches in this study (Supplementary Fig. 6a-d) and across our previous work using this cell line ^42,43^. RPE-1 showed a diploid karyotype (Supplementary Fig. 6c-d), with a subset of metaphases showing 42-45 chromosomes likely due to sliding during metaphase preparation (Methods), and no evidence of polyploidy, tetraploid or pseudo-tetraploid clones in all samples analyzed (Supplementary Fig. 6a-d).

Furthermore, no gross chromosomal rearrangements (GCRs) or other cytologically visible chromosomal changes were observed through passages and comparing batches of RPE-1 acquired from different sources, implying that RPE-1 cells have a remarkably stable diploid asset maintained in culture (see “Discussion”). Altogether, the resulting final RPE-1 genome (RPE1v1.0) represents the first haplotype-resolved ultra-high quality complete reference of an experimentally-amenable diploid human cell line.

### Comparative characterization of RPE-1 inter-genomes and inter-haplotype variation

Recent analyses of a completely phased diploid genome suggest that differences between the two haplotypes of the same individual carry variation within coding regions ^2^ that increase in polymorphic loci like centromeres which limit meiotic crossover ^47,48^. To establish RPE-1 specific variation, we characterized the specific genomic make-up of RPE-1 for each phased haplotype (Fig. 2a). First, we wanted to define the haplotype with the translocated X and map the precise breakpoint in the t(Xq;10q) at base pair resolution. A previous attempt determining chromosomal haplotypes inferred by bulk DNA sequencing reported high likelihood of the RPE-1 translocated segments belonging to the same haplotype ^46^. We instead developed a multi-step pipeline for the manual curation of the Structural Variant (SV) starting from the fully phased haplotypes (Fig. 2b, Supplementary Fig. 7). With this approach (Supplementary Note 3), we identified a 73.18 Mb segmental duplication of chromosome 10 *q-*arm most likely to belong to Hap 2 (due to read alignment score) (Supplementary Fig. 7a), fused to the telomeric region of Hap 1’s chromosome X. Validation of the breakpoint also identified an ensuing deletion of 3603 bp mapping proximally to the X’s telomere and associated with the chromosomal fusion (Fig. 2b, Supplementary Fig. 7b-c).

**Figure 2.**
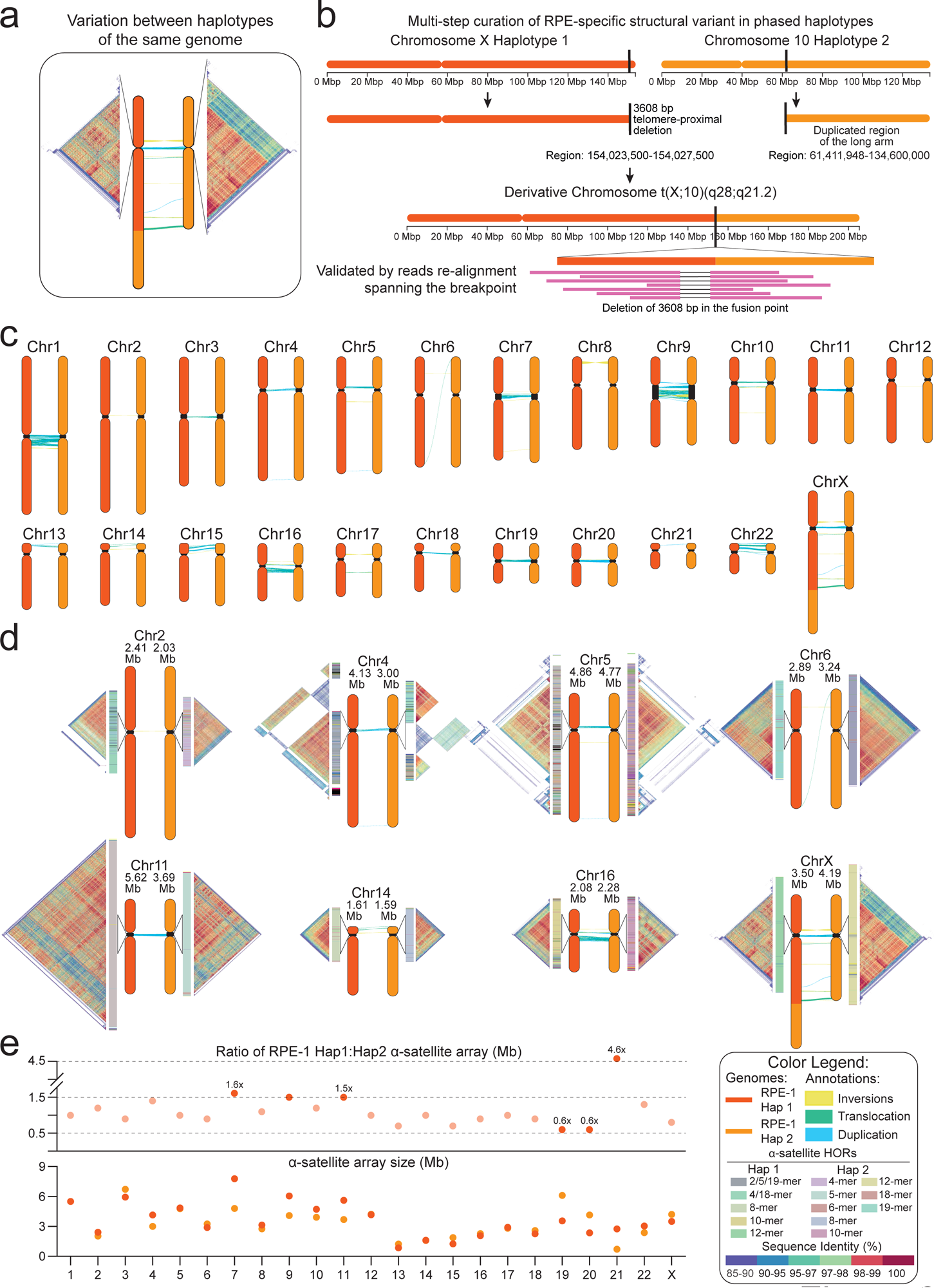
Genome variation between RPE-1 haplotypes. (a) RPE-1 haplotypes harbor repeat structure variations at centromeres, as shown by the pairwise similarity heatmaps of the HORs (Methods). (b) Base-level resolution of *t*(X;10) (*q*28;*q*21.2), a RPE-1-specific structural variant (Methods) involving a 73.18 Mb segment, duplicated from the *q*-arm of Hap 2’s chromosome 10 fused to the microdeleted telomeric region of Hap 1’s chromosome X. (c) Synteny between RPE-1 haplotypes highlight intra-haplotype structural variations, peaking at centromeric and pericentromeric regions. (d) Pairwise similarity heatmaps of chromosomes with major intra-haplotype differences in term of centromere sequence, size, structure and position. HORs structural variants (SVs) are shown for each chromosome. (e) Ratio (top) of centromere length (bottom) between the two RPE-1 haplotypes. Chromosomes with a length ratio greater than 1.5 or less than 0.7 are highlighted.

Next, we assessed genome variation between the two RPE-1 haplotypes and between each haplotype and the CHM13 using SyRI ^49^. As expected, we observed that RPE-1 and CHM13 are syntenic throughout most of the genome with 60% of whole-genome variants falling within highly diverged regions (HDRs), repetitive and polymorphic loci (Fig. 1c, Table 2). Similarly, RPE-1 haplotypes showed true-positive variation rates including a total of 62 inversions, 1646 translocations and 2565 HDRs, with the greatest differences occurring in the centromeric and pericentromeric regions (∼40%) (Fig. 2c). Notably, both RPE-1 haplotypes carried a similar number of SVs when compared against CHM13: 50 inversions, 876 translocations and 2589 HDRs for RPE-1 Hap 1, and 76 inversions 1602 translocations and 2636 HDRs for Hap 2 (Table 2) associated with the Major Histocompatibility Complex (MHC), one of the most gene-dense and polymorphic stretches of human DNA ^50^ and centromeric and pericentromeric regions (Fig. 2c, Table 2).

A resolved diploid genome gave us an opportunity to investigate intra-individual variation between the two haplotypes of RPE-1 for the first time (inter-haplotype differences, Fig. 2a). To do so, we focus on the most polymorphic regions and annotated centromeres with HumAS-HMMER_for_ANvil and RepeatMasker for all chromosomes ^51^ (Methods). We found that centromere alpha-satellite arrays in RPE-1 range from 0.4 (chromosome 21, Hap 2) to 7 Mb (chromosome 7, Hap 1) (Fig. 2d-e), a wider size range than previously found for other cell lines. Given previously reported recombination suppression in these regions ^47^, we anticipated observing substantial variation between centromeres of homologous chromosomes and likely, haplotypes. In line with this, RPE-1 centromeres of the same chromosome pairs show large differences not limited to their size (Fig. 2d-e) but also HOR structure inferred by pairwise sequence identity heatmaps (Fig. 2d) using StainedGlass (v6.7.0) ^28^ (Methods). Indeed, we found inter-haplotype differences in the centromeres of all chromosomes, with 49% of them showing variations in organization, number of monomers in the HOR, specific HORs SVs and length within the active centromere (Fig. 2d, Supplementary Fig. 8). For example, inter-haplotype structural divergence was observed for chromosome 11, 22 and X, while chromosome 6 of Hap 1 also presented a unique Live Active HOR (LHOR) with two highly homozygous regions (Fig. 2d). Similarly, chromosome 4 of Hap 2 and chromosome 19 of Hap 1 showed several repetitions of the same monomer motif along the HOR region suggesting the insurgence of new active HORs (Fig. 2d, Supplementary Fig. 8, Table 3), likely through previously suggested layered expansion ^22^. These observations underscore that, structurally, centromere of one haplotype may be anchored within a specific type of single live HORs while the other relies on a different or expanded structure, reminiscent of metastable epialleles that had been cytologically shown for chromosome 17 ^52^.

Comparison with HG002v0.7 showed similar inter-haplotype variation for all chromosomes, indicating that our finding may be widely applicable to human centromere polymorphism in diploid genomes. Notably, previously reported centromeric inversion on chromosome 1 HORs of CHM13 ^1,22^ was not found in either RPE-1 or HG002 haplotypes ^2^, suggesting that it may represent a rare or specific polymorphism of that line ^22^. However, we found smaller haplotype-specific inversions ^30^ within the centromeres of RPE-1 and HG002, for chromosomes 5 and 6 for the maternal genome and for chromosome 3 of Hap 2. Next, we evaluated RPE-1 inter-haplotype variation by calculating the ratio of centromere sizes. We found a distribution between 0.5- and 1.5-fold differences, with chromosomes 7, 11 and 21 having the greatest size divergence between haplotypes (Fig. 2e). Interestingly, the centromere of chromosomes 7 and 21 showed more than 1.5 increase in size when comparing the RPE-1 haplotypes, suggesting that they may carry higher divergence or be more challenging to assemble and validate (Fig. 2e). This finding also raises the question of the biological impact of size divergence. For chromosome 21, this has been associated with ensuing chromosome missegregation during embryonic development that may underlie Down’s Syndrome ^53^; for chromosome 7, biological relevance remains to be established. Altogether, our data present evidence and quantify the extent of the polymorphism between haplotypes of the same genome.

### Alignment improves using RPE-1 genome compared to non-matched references

The haplotype variation and divergence between genomes found using the RPE-1 reference is case-in-point to support the need for matching the reads generated from multi-omics experiments with the reference genome used to align them. Non-aligned content to widely-used human reference HG38 has been shown in all human genomes analyzed ^54^. While for small genomes it has been possible to study genomic variation by whole-genomes comparison ^55,56^ for large human genomes, matched sequence-reference has been previously considered but remained largely unexplored ^57^. Recent efforts addressed reference bias and its negative impact on read mapping by increasing representation of individuals’ genomes variation ^3,18,58^. We reasoned that alignment and analyses of multi-omics data may not always be successfully supported using a non-matched single reference, or a collection of them, particularly for regions that show considerable variation (Fig. 2c-d). To validate the improvement in read alignment when a matched reference genome is used, we aligned RPE-1 HiFi reads (∼46x) against the RPE-1 diploid genome using the complete CHM13 assembly as control (Fig. 3). Because CHM13 is a haploid genome, we chose a different approach to show not only the differences between completely matched and non-matched genomes, but also to highlight the different alignment between diploid and haploid genomes. (Fig. 3a). Long read alignment was performed using Minimap2 (Methods), without applying any quality or mapping filters, as the use of them led to the loss of the reads mapping in the homozygous regions of the diploid genome (∼80%). We observed uniform coverage on both haplotypes compared to the expected 20x coverage, as opposed to CHM13 that showed relatively uniform alignment coverage but with higher mismatched bases. Diploid alignment of HiFi reads is equally distributed between each RPE-1 haplotype (20x) but secondary alignment and multimapping was much higher mapping to the CHM13 genome, further increasing within repetitive regions (Fig. 3b). We then tested the improvements in quality and fidelity of alignment of isogenomic RPE-1 reference and CHM13 on centromeric regions for chromosome 1, 2, 4, 5, 6, 11, 16 which presented the highest diversity in read coverage between these two cell lines (Fig. 3b, Supplementary Fig. 9). We observed a lower coverage or absence of mapped reads for the CHM13 alignment, while a complete uniform coverage for the RPE-1 (Fig. 3b, Supplementary Fig. 9a). This result is explicable by the high divergence in sequence identity for centromeric regions comparing these two genomes (Fig. 3b, Supplementary Fig. 9b). These data point to an improvement in coverage, quality, fidelity and, consequently, a reduction in incorrectly aligned reads upon use of the isogenomic reference for sequencing alignment.

**Figure 3.**
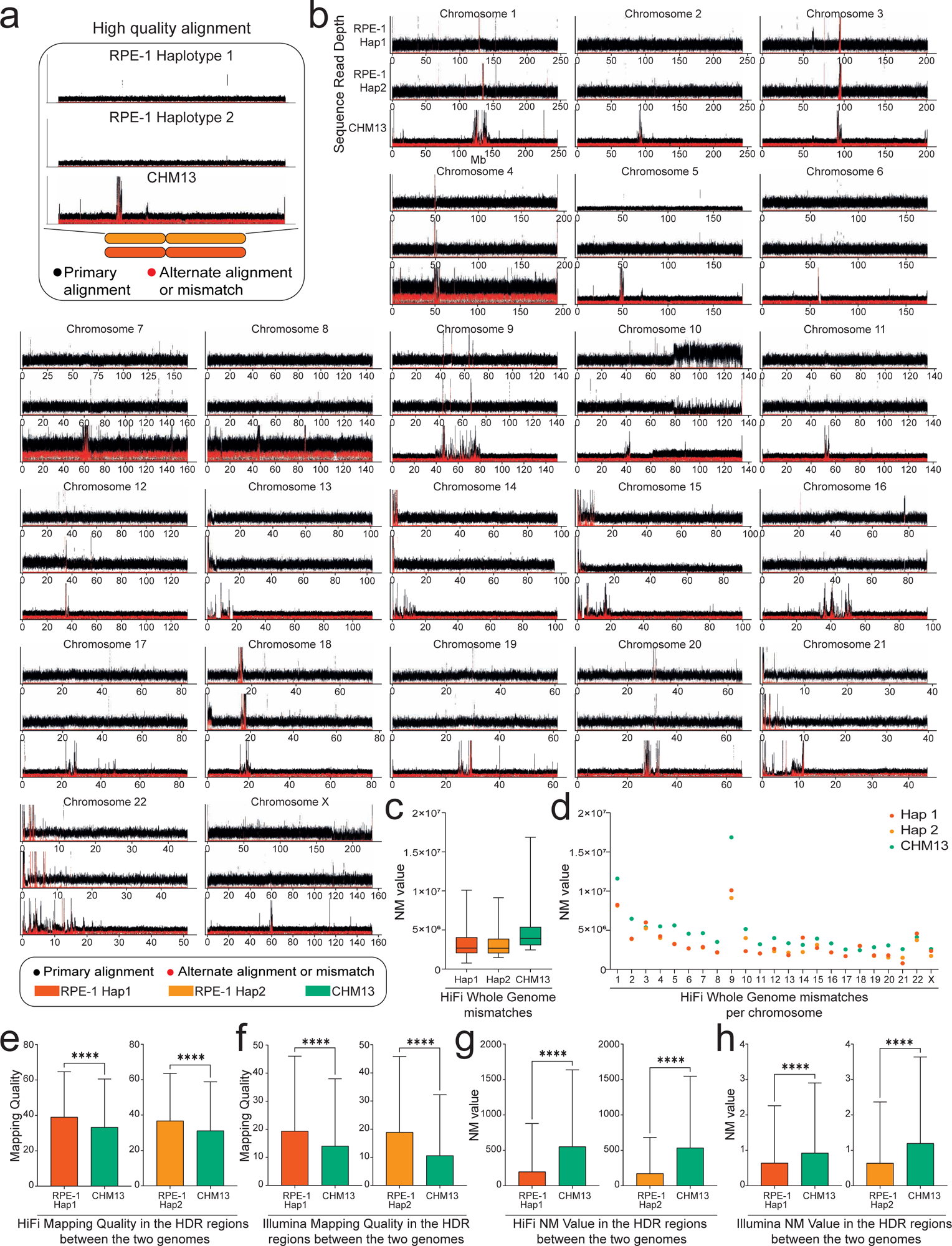
The isogenomic reference genome improves reads alignment. (a) Read-depth profiles over RPE-1 Hap 1, RPE-1 Hap 2, and CHM13. The black and red dots represent, respectively, the coverage of the first and second most frequent alleles. (b) HiFi read-depth profile for all chromosomes. Using the RPE-1 haplotypes as reference improves the primary alignments compared to CHM13. The differences in coverage for long arm of chromosome 10 in RPE-1 genome are due to regional duplication (Methods). (c) Sequencing reads from a different batch of RPE-1 show lower edit distance (NM value) when aligned against the matched reference compared to CHM13, consistent across all chromosomes (d). HiFi and Illumina reads from High Diverged Regions (HDRs) show statistically significant increase in mapping quality (e-f), and lower NM value (g-h), when aligned against the RPE-1 haplotypes compared to other reference. p-values are from the student *t*-test (non-significant (ns) = p > 0.05, * p ≤ 0.05, ** p < 0.01, *** p < 0.001, and **** p < 0.0001).

To confirm and validate these improvements, we statistically evaluated the differences in read alignment using two main values: 1) mapping quality and 2) edit distance to the reference, defined as the number of changes necessary to make an aligned read equal to the reference, excluding clipping (NM value) (Supplementary Note 4). This analysis was performed using whole-genome sequencing (WGS) reads generated with a different batch of RPE-1 cells, sequenced with two different technologies: Illumina short reads and HiFi long reads. We mapped these reads using BWA and Minimap2 (Methods) against Hap 1, Hap 2 and CHM13, without using the final diploid genome and applying quality filters, excluding secondary, supplementary and multimapped reads during the alignment step. Mapping quality and NM value were evaluated for each read which belongs to the final alignment, comparing the distribution of both values for Hap 1, Hap 2 and CHM13 genomes. Whole genome alignment showed a decrease in NM value using RPE-1 haplotypes as a reference (Fig. 3c, Supplementary Fig. 10a-b) and an average distribution of mapping quality around score of 60 (which represents the highest value in mapping quality scores, highlighting the reliability of the mapped read) for all genomes (Supplementary Fig. 10c-d). Despite mapping quality of Hap 1 showed mostly a distribution around score of 60, we observed a statistically significant lower score compared to CHM13 (p-value <0.0001 evaluated by student t-test) (Supplementary Fig. 10d). To explain this result, we then evaluated mapping quality and NM value for each chromosome separately (Fig. 3d, Supplementary Fig. 10e). In the marker chromosome (translocated chromosome X) of Hap 1 we observed a decrease in both the NM value (Fig. 3d) and the mapping quality score (Supplementary Fig. 10e), thus confirming that such lower values were determined by the presence of the duplicated long arm of chromosome 10, and not caused by the presence of misassembled regions for this chromosome.

We then focused on syntenic HDRs, the most diverged between RPE-1 and CHM13. HDRs were extracted from the output of SyRI, obtaining a total of 37% and 57% (in bp) of HDRs belonging to LHOR when Hap 1 and Hap 2 were compared to CHM13 (Supplementary Fig. 11). Our *de novo* assembly did not show any regional or structural errors at these regions, ensuing the possibility to completely compare RPE-1 to CHM13 (Supplementary Fig. 11, Table 2). Within the HDRs, both mapping quality score and NM value significantly changed when RPE-1 was used instead of CHM13, with a ratio of 3.5 for mapping quality score (comparing Hap 1/Hap 2 on CHM13) (Fig. 3e-f) and a 2.1 for the NM value (comparing CHM13 on Hap 1/Hap 2) (Fig. 3g-h). In particular, the HDR regions identified between Hap 2 and CHM13 showed a lower mean distribution of mapping quality score (Fig. 3e-f) and higher NM value (Fig. 3g-h) for CHM13 compared to Hap 2. Indeed, Hap 2 showed the lowest NM value, confirming the presence of peculiar sequences in its polymorphic loci, as highlighted by the higher number of SVs found using the syntenic region founder tool (Methods). The same level of accuracy is not supported by the latest complete human reference genome CHM13, in spite of being the most complete and validated genome available to date. Altogether, our results showed that the reference bias decreases while uniquely mapped reads increase when using a matched reference, carrying out an innovative resource in SVs analysis phasing evaluation in protein DNA interaction experiments ^59^.

### Isogenomic reference enables correct epigenetic characterization of human centromeres

To verify whether the isogenomic reference genome can improve the analysis of epigenetic data in highly polymorphic regions, we used a publicly available CUT&RUN dataset from RPE-1 cells and we compared the results by using different reference genomes: RPE-1 haplotypes (RPE1v1.0 Hap 1 and Hap 2), CHM13, HG38 (GRCh38.p14) and the HG002 haplotypes (HG002v1.0 paternal and maternal). We focused on the analysis of centromeric chromatin enrichment and spread of the centromere-specific H3 histone variant, CENP-A. Recent studies have shown differential enrichment loci during the cell cycle ^60^, changes under different treatment conditions ^61^ and, more recently, shifts in the region of CENP-A enrichment called through non-isogenomic alignment ^62^. We used the RPE-1 reference to re-assess and improve multi-omics analyses of human centromeres by aligning a CUT&RUN a publicly available dataset (GSE132193) ^63^ (Methods) performed on RPE-1 cells (Fig. 4a). Since the epigenetic centromere identity is defined and maintained by the presence of the histone variant CENP-A, we used experiments conducted on RPE-1 to explore the power of isogenomic reference genome for the investigation of the structure and organization of the centromeres at higher level of epigenetic precision due to improved reads mapping (Fig. 3). Mapping of CENP-A-bound reads show dramatic differences in occupancy and spread at nearly all centromeres between the RPE-1 haplotypes and HG38 (Fig. 4a-d, Supplementary Fig. 14). This is in line with lack of complete annotation, HG38 shows incomplete peak calling on chromosomes 1, 9 and 16 which all have a large array of HSat2 and HSat3 ^1^, likely due to incomplete centromeres’ organization and annotation in the HG38 genome which impacts the ability to distinguish between centromeric and pericentromeric regions. However, even using CHM13, which has been used to study the epigenetic landscape of these loci ^62^ and contains complete linear sequence for all centromeres ^1^, CENP-A enrichment is not assigned correctly compared to peaks calling alignment on RPE1v1.0 (Fig. 4a-d). The variation in the position, size and shape of peaks of the centromere marker proteins between CHM13 and RPE-1 is likely due to the ability to allow reads originated from a diploid cell line to map on the phased haplotypes to better determine a *bona fide* primary alignment. This is especially significant for polymorphic and repetitive regions of the human genome where multimappers can present more than one possible alignment site, indicating that having the matched reference unlocks increased accuracy in the discernment of the most likely alignment site. To address whether the increased accuracy was truly dictated by the isogenomic references instead of the mapping on a diploid genome, we extended our comparison to the only other currently available complete diploid assembly of HG002 ^1,2,64^. As we found for CHM13, the enrichment of CENP-A changed based on the haplotype used but was never comparable to using RPE-1 genome as its own reference (Fig. 4d). Next, we wanted to determine if the epigenetic data mapping RPE1 reads isogenomically were indeed “the correct alignments”. To do so, we investigated the structure of the active centromere by monitoring the presence and the spread of CENP-A enrichment *via* high confidence CENP-A peaks with a ueue ≤0.00001 (Fig. 4b, Methods). We found that the use of matched reads-reference was the only condition that enabled the detection of a single cluster of CENP-A peaks, while the high confidence peaks called on all other genomes analyzed were scattered along the active centromere array (Fig. 4a-b). We reasoned that this precise mapping of CENP-A enrichment may represent the exact positioning of the kinetochore (Fig. 4b). To validate if we were able to harness the isogenomic reference genome to determine the exact kinetochore site at an unprecedented precision, we confirmed the co-localization of the high-confidence CENP-A peaks (MAPQ >20, q-value ≤0.00001) with the cytosine methylation (5mC) positioning that identifies the hypomethylated CDR (Methods). The CDR has been recently highlighted as the site predictive of the functional region within the active centromere for the binding of the kinetochore ^22^. We found colocalization between high confidence peaks and the CDR (Fig. 4b-c). Notably, we were able to map the precise site within the region marked by the drop in methylation at base pair resolution, unveiling the sequence and structure of the centromere HOR underlying the kinetochore which differs within haplotypes (Fig. 4b), further underscoring the need to use phased genomes for epigenetic precision at divergent loci. The centromere of chromosome 2 shows the lowest variation in kinetochore position between the two RPE-1 haplotypes and, interestingly, exhibits the lowest variation in size for the CDR (Fig. 4b). Moreover, chromosome 2 displayed the highest conservation of CENP-A mapping among the reference genomes, and this is further reflected by the high conservation in size of the active centromere (Fig. 4b). On the contrary, chromosome 7 presented the highest, near-2-fold expansion, in CENP-A occupancy both regarding RPE-1 haplotypes and to HG002 haplotypes, highlighting an underlying variation at the sequence level that, in turn, changes the epigenetic landscape of the centromere (Fig. 4a, Supplementary Fig. 14). Surprisingly, when assessing the CDRs, we noticed that these differences in chromosome 7 are not reflected in the kinetochore’s size of the RPE-1 haplotypes (Fig. 4a). The centromeres of chromosome X and 7 both exhibited an interesting 440-500 kb respective range of distance between the CDRs of the two haplotypes (Supplementary Fig. 14). One can appreciate the power of the isogenomic referencing especially when focusing on the reliable detection and calling of the CDRs which would have been missed otherwise because it falls within a region which would have been hidden when mapping against the other assemblies. Also, the absence of the underlying sequence and structural expansion in other haplotypes would have made it impossible to identify this enrichment without matched-reference. Importantly, the alignment between the high-confidence CENP-A peak and the CDR would be shifted using a reference genome in all cases observed (Fig. 4a-c). The precision of mapping also enabled us to call enrichment spot for CENP-A identifying the site of the kinetochore to be occasionally localized toward the left or right boundary of the CDR (Fig. 4c), implying a structural organization that may favor chromosome segregation (see “Discussion”). Interestingly, our reference showed that location of the CENP-A high-confidence peaks did not have a preferred positioning within the CDR – i.e. either toward the left or right side of the dip (Fig. 4b-c, Supplementary Fig. 14), suggesting a minimal requirement of CENP-A nucleosomes to be productive for kinetochore assembly and chromosome segregation. Altogether, we have highlighted a significant variation between cell lines that affects downstream analysis and interpretation, showing the power and need of matched reference genome for multi-omics analyses.

**Figure 4.**
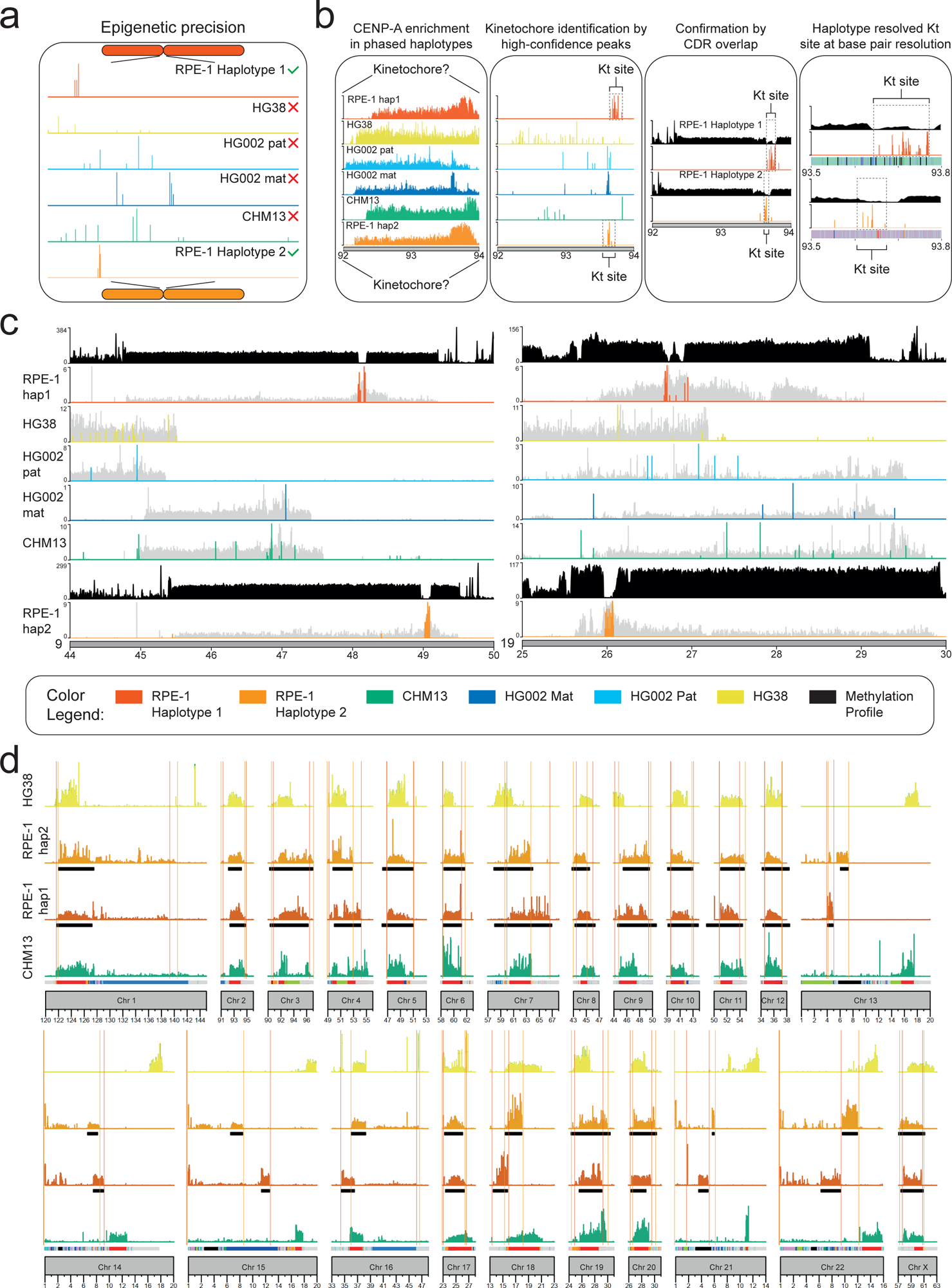
The isogenomic reference genome identifies high-confidence epigenomic landscapes. (a) Phased peak calling improves when using the isogenomic genome as reference. (b) Pipeline describing the mapping of CENP-A CUT&RUN enrichment to reveals the precise kinetochore (KT) position with high confidence. High confidence CENP-A peak map to a single genomic location that differs between haplotypes defining the kinetochore site only when the isogenomic reference is used but not with any other reference tested, giving scattered or low significance peaks. This is confirmed by methylation analysis to overlap with the CDR. Kinetochore sequence and structure can be defined at base pairs resolution. (c) Example of methylation track (defining the CDR) and CENP-A high-confidence peaks (defining the kinetochore) for chromosomes 9 and 19 only using RPE-1 as reference but not with CHM13, HG38, HG002 mat or pat genomes. (d) The spread of CENP-A occupancy changes between haplotypes for all chromosomes, and when using different genomes raising questions on the validity of non-matched references for mapping reads to such polymorphic loci.

## DISCUSSION

Here we present the first, to our knowledge, human reference genome of a diploid experimentally-relevant laboratory cell line, including fully-phased haplotypes spanning the centromeres, RPE-1v1.0. We found four differences at human centromeres: (1) in sequence, (2) in size, (3) in structure, and (4) in position. In line with rapid emergence of new alpha-satellite HOR structure that underlies divergence in centromeres between genomes ^31^, we found differences in one or more of these 4 parameters between haplotypes for all chromosomes. With this extent of divergence in mind, we demonstrate how the use of this *isogenomic* reference genome better supports accurate multi-omics analyses of reads derived from RPE-1 cell line, especially for regions with the highest diversity, compared to the common practice of using the available human reference genomes. We show that RPE-1 reads from centromeric chromatin map differently depending on the reference genome used without necessarily reflecting a *bona fide* biological change. This is noteworthy considering the abundant literature showing epigenetic datasets calling CENP-A localization ^42,60–62,65–67^, and other centromeric proteins ^43,68,69^, previously mapped on reference genomes. Isogenomic mapping using RPE-1 improves all alignments parameters tested, using whole-genome or HDR and either long or short reads compared to the latest references, CHM13 and HG002 maternal and paternal haplotypes ^1,2,37^ and HG38. Not only isogenomic mapping substantially improves peaks positioning, size and enrichment confidence compared to all other genomes but also enables to identify high-precision phased epigenomic landscape that adds valuable information to interpret the findings in a diploid state. Thus, our work addresses the reference bias ^18^ empirically demonstrating how sequence changes negatively impact the ability to define, size and position of the active centromeric chromatin. To this end, we show for the first time the precise kinetochore site within the CDR only using the isogenomic reference RPE-1v1.0 to align RPE-1 reads from CENP-A chromatin immunoprecipitation experiments but not using any other reference genome – however complete. Reanalyzing isogenomically previously published datasets ^62^, we found variation in kinetochore position between haplotypes in all chromosomes. These remarkable inter-haplotype differences are in line with recent observations of high kinetochore plasticity between genomes ^31^. We determined the enrichment ratio for chromosomes 2, 7 and X which shows higher, equal or lower CENP-A levels between haplotypes, underscoring this heterogeneity. Interestingly, the CDR ratio showed lower levels of variation between haplotypes, suggesting that isogenomic mapping identifies the minimal functional size required, often positioned proximal to the beginning or end of the CDR. CENP-A has been previously shown to be peppered along the active HORs, and its density increases from an estimate of 1 every 20 to every 4 canonical nucleosomes. Our data imply that the such CENP-A enrichment positioned at the CDR boundary – left or right, marks the exact sites of kinetochore attachment for that haplotype of each specific chromosome. Having this level of haplotypes-resolved sequence-specific precision will transform technologies like genome editing and chromosome-specific aneuploidy ^70^.

Only recently, the enduring quality of HG38 assembly patchwork was celebrated as a valuable reference ^71^; today, we demonstrate a conceptual shift to address the reference biases and sequence variation ^72^ using matched reads-reference. Thus, while the human Pangenome draft ^3^ represents an important step toward a more equitable and fair representation of human genomic diversity ^3^, our work expands from the idea of representing as many human variants as possible toward the concept of aligning to a “personalized” reference sequence that already incorporates the relevant individual’s variants ^57^. We selected RPE-1, because, besides being widely used across fields with over 2000 publications ^35^, their stable karyotype offers a reference genome that can in principle support experimental data generated in any laboratory.

Altogether, our study opens important opportunities pertaining to advancing our understanding of the extent of the reference bias, its influence on downstream analysis and the biological relevance of improved alignment for genomic and epigenomic studies of genomes, especially at HDR such as centromeres. Our proof-of-concept calls for a comprehensive catalog of complete genome assemblies for commonly used cell lines for a widespread application of *isogenomic* reference genomes for faithful multi-omics analysis changing downstream analyses, interpretations and discoveries. Our aim is to build a comprehensive catalog of complete genome assemblies for commonly used cell lines, including diploid embryonic and induced pluripotent stem cells (ESC and iPSC), primary and cancer cell lines. Finally, integrating the human pangenome graph that represents human diversity with genome assemblies derived from historic experimentally-amenable cell will provide important information about the differences between reference genomes freshly derived from individuals and those generated from experimentally-amenable laboratory cell lines. The RPE-1 genome represents a new telomere-to-telomere human diploid reference for the scientific community that will advance genetic and epigenetic research across fields using this cell line and a first step toward “personalized” genomes – for the benefit of individuals and multi-omics studies alike.

## METHODS

### Cell lines

The hTERT-immortalized retinal pigment epithelial (hTERT RPE-1) cells used to generate the RPE-1v1.0 assembly were validated by MSKCC; a second batch of cells was purchased from ATCC (CRL-4000); a third batch of cells was obtained from a laboratory in a different continent ^70^. RPE-1 cells are near-diploid human cells of female origin with 46 chromosomes. Cells were cultured in DMEM/F12 medium supplemented with 10% fetal bovine serum (Gibco 10270-106), 100 µg/ml streptomycin, 100 U/ml penicillin (Euroclone ECB3001D) and 2 mM glutamine (Euroclone ECB3000D). Cells were grown at 37°C in a humidified atmosphere of 5% CO_2_, and tested negative for mycoplasma contamination.

### Metaphase spread preparation

hTERT RPE-1 cells at 70% confluence were harvested by trypsinization after 2-hour treatment with colcemid (100 ng/ml, Roche 10295892001) to arrest cells in mitosis, washed with PBS and incubated with a pre-warmed hypotonic solution (0.07 M KCl) for 30 minutes at 37°C. After centrifugation, swollen cells were fixed with a solution of methanol-acetic acid (3:1) and washed twice with the same fixative solution. Fixed cells were dropped onto clean, wet slides and air dried overnight. All centrifugation steps were performed at 1,200 rpm for 5 minutes at room temperature (RT).

### Karyotype analysis by R-banding with Chromomycin A3

To confirm the chromosomal structure of the assembly, a karyotype for hTERT RPE-1 cells was generated. Metaphase spreads from hTERT RPE-1 cells were washed with a phosphate buffer (0.07 M NaH_2_PO_4_, 0.07 M Na_2_HPO_4_, 1 mM NaCl, pH 6.8) followed by 2-hour incubation with Chromomycin A3 (0.6 mg/ml, Sigma-Aldrich C2659) at RT in a dark, moist chamber. Slides were then washed with NaCl-HEPES (4-(2-hydroxyethyl)-1-piperazineethanesulfonic acid) (0.15 M NaCl, 5 mM HEPES), stained for 15 minutes in a methyl green solution (0.1 mM, Sigma-Aldrich) and washed twice with NaCl-HEPES. The antifading (Vectashield H-1000; Vector Laboratories) isopropilgallate 1:300 was added to the slides before storing them at 4°C for 3 days in a dark, moist chamber. Images were acquired using a Thunder fluorescent widefield microscope (Leica) at a 100X magnification.

### DNA extraction

hTERT RPE-1 cells at 70-80% confluence were harvested by trypsinization, washed with PBS and centrifuged at 1,000 rpm for 5 minutes at RT. Cell pellets were prepared in individual aliquots of 1.5 x 10^6^ cells, frozen in dry ice and stored at -80°C until further use. High Molecular Weight (HMW) DNA and ultra-high molecular weight DNA (UHMW) was extracted from hTERT RPE-1 dry cell pellets using the Monarch HMW DNA extraction kit for cells & blood (New England Biolabs, NEB T3050L) and for tissue (NEB T3060L), following manufacturer’s instructions and introducing few technical improvements. HMW DNA was quantified by Nanodrop and Qubit with a broad range kit (Thermo Scientific Q32850). Native DNA size distribution assessed using Femto Pulse with Genomic DNA 165 kb kit (Agilent FP-1002-0275).

### Library preparation and sequencing

Sequencing data were obtained using two complementary long-read sequencing technologies for the assembly of hTERT RPE-1 cells: Pacific Biosciences (PacBio) High-Fidelity (HiFi) long reads and Oxford Nanopore Technologies (ONT) long (>70 kb) and Ultra-Long (UL; >100 kb) reads. Additionally, Illumina and Hi-C (Arima Genomics) sequencing technologies were used.

HMW DNA from 1.5 x 10^6^ cells was used to generate PacBio HiFi libraries with the SMRTbell express template prep kit 2.0 (PacBio 100-938-900). Size selection was performed with Megaruptor 2 (Diagenode) with default settings, and fraction sized 15-20 kb as determined by Femto Pulse were sequenced on the Sequel IIe platform with SMRT Cells 8M (PacBio 101-389-001) using the chemistry 2.0 with 2-hour pre-extension, 2-hour adaptive loading and 30-hour movie collection time, to reach a coverage of 46x in PacBio HiFi reads. Circular Consensus Sequencing (CCS) reads were obtained using SMRT-Link (https://github.com/WenchaoLin/SMRT-Link) with default parameters.

UHMW DNA from 6 x 10^6^ cells was used to generate UL-ONT libraries with the UL DNA sequencing kit (ONT SQK-ULK114) following manufacturer’s instructions. Ninety µl of library was loaded in a R10.4.1 (FLO-PRO114M) flow cell and sequenced on the PromethION 24, with two nucleases washes and reloads after 24 and 48 hours of sequencing to reach a total coverage of 80x in ONT reads (average 70 kb) and of ∼30x in 100 kb + ONT reads with a Q score >20. These ONT data were base-called with Guppy (v5.0.11). In parallel, HMW DNA from 6 x 10^6^ cells was used to generate additional UL-ONT libraries with the same UL DNA sequencing kit and protocol. Ninety µl of library was loaded in a R10.4.1 (FLO-PRO114M) flow cell and sequenced on the PromethION 2 Solo, with two nucleases washes and reloads after 24 and 48 hours of sequencing to reach a total coverage of 10x in ONT reads (average 19 kb) with a Q score >35. These ONT data were base called with Guppy (v6.5.7).

HMW DNA from 1.5 x 10^6^ cells was used to generate Illumina libraries with the Nextera XT DNA Library Preparation kit (Illumina FC-131-1024) and Illumina DNA PCR-Free Library Prep, Tagmentation Library Preparation kit (Illumina 20041795) following manufacturer’s instructions. These libraries were sequenced on the Illumina Nova-seq 6000 to reach a coverage of 120x and 60x in Illumina reads. Raw data were processed with Fastp (https://github.com/OpenGene/fastp) ^73^ to trim the adapters and FastQC (https://github.com/s-andrews/FastQC) ^74^ to evaluate the quality of the reads.

HMW DNA from 2 x 10^6^ cells was used to generate Hi-C libraries with the Arima High Coverage HiC+ kit (Arima Genomics A101030), and the Arima Library Prep Module kit v2 (Arima Genomics A303011) according to the manufacturer’s protocols. Hi-C libraries were sequenced on the Illumina Nova-Seq 6000 to reach a coverage of ∼30x in Hi-C reads. Raw data were processed with Cutadapt (https://github.com/marcelm/cutadapt) ^75^ to trim the adapters.

### Whole genome assembly, manual curation and phasing pipelines

Before assembling the genome of hTERT RPE-1 cells, global genomic features such as heterozygosity, repeat content and size were evaluated with GenomeScope 2.0 (https://github.com/tbenavi1/genomescope2.0) ^76^ from unassembled HiFi raw sequencing reads.

Two different assembly algorithms were then employed, Hifiasm (https://github.com/chhylp123/hifiasm) ^12,36^ and Verkko (https://github.com/marbl/verkko) ^37^. The assembly was first generated with Hifiasm using HiFi data with base-calling accuracy of 99.99%. This first assembly was compared to the assembly generated with Verkko both using HiFi together with ONT UL reads only, and using HiFi and all ONT reads regardless of size, with integration of Hi-C linkage data for manual curation and complete haplotypes phasing of the human diploid genome. The VGP mapping pipeline ^77^ was run to map the Hi-C reads against Hap 1 and 2 independently and against the diploid assembly. The final diploid assembly.fasta was merged with the unassigned.reads.fasta generated by Verkko. On PretextView (https://github.com/wtsi-hpag/PretextView), the contact map derived from the conversion of the last alignment file using PretextMap (https://github.com/wtsi-hpag/PretextMap) was used, and the correct position was assigned for each contig. The subsequently dual manual curation was done as described in Rapid-curation-2.0 (https://github.com/Nadolina/Rapid-curation-2.0) achieving 23 chromosome-level scaffolds for each haplotype. The algorithm MashMap (https://github.com/marbl/MashMap) ^78^ was then used to perform a genome-to-genome alignment, with the CHM13 assembly as a reference, in order to identify each chromosome. MUMmer (v4.x) (https://github.com/mummer4/mummer) ^79^ and GSAlign (https://github.com/hsinnan75/GSAlign) ^80^ were used to make dot plots of each chromosome and determine whether the orientation of the chromosomes in the assembly was correct.

### Evaluation of de novo genome assembly

To assess the quality of the final RPE-1v1.0 assembly, reference-based assemblers such as Cross-stitch and MaSuRCA were purposely avoided, to obtain an unbiased genome evaluation completeness and quality without using a genome comparison. The quality and gene completeness were evaluated using the reference-free Merqury *k*-mer analysis tool (https://github.com/marbl/merqury) ^38^, BUSCO (https://github.com/WenchaoLin/BUSCO-Mod) ^39^ and Asmgene ^40^ with default parameters. The Bandage (https://github.com/rrwick/Bandage) ^81^ visualization on the string graph ^82^ is used to represent continuous whole-chromosome contigs and unassigned reads. The quality of the assembled genome was evaluated with Gfastats (https://github.com/vgl-hub/gfastats) ^83^ and QUAST (https://github.com/ablab/quast) ^84^. We assessed the assemblies for uniform read depth across chromosomes via IGV ^85^ and NucFreq (https://github.com/mrvollger/NucFreq) as well as read mapping bias. Assembly errors were evaluated using Craq algorithm, which uses clipping information mapping the original sequencing reads back to the de novo genome assembly, and defining a reference level Assembly Quality Indicators (AQI over 99%) ^41^ (Fig. 1b, Table 2).

The read-based pipeline Flagger ^3^ was used to detect different types of misassemblies within a phased diploid assembly.

### Identification of Structural Variants

Minimap2 (https://github.com/lh3/minimap2) ^40,86^ performed a genome-to-genome alignment choosing the parameter -x asm5 for genomes with low sequence divergence. SyRI (https://github.com/schneebergerlab/SyRI) ^49^ was used to search variants between: Hap 1 vs. Hap 2 of the RPE1v.1.0 assembly; CHM13 vs. Hap 1; and CHM13 vs. Hap 2 with default parameters. SyRI outputs the complete information about Structural Variants and genomic rearrangements, such as syntenic regions, copy number variation and highly diverged regions between two or more genomes. Final visualization was obtained with Plotsr (https://github.com/schneebergerlab/plotsr) ^87^.

### Multi-step identification and curation of breakpoint in phased haplotypes

To map the precise t(Xq;10q) breaking point characteristic of hTERT RPE-1 cells ^35^, we performed a stepwise process (1) detecting changes in read coverage for both haplotypes; (2) read alignment on the diploid genome to identify haplotypes with translocated X; (3) isolation of reads spanning the breakpoint on chromosome X, (4) manual curation of breakpoint at base pair resolution, and (5) validation of the new fasta file through reads re-alignment. HiFi and ONT read alignments were first evaluated against Hap 1 and 2 of our *de novo* RPE-1 genome (RPE-1v1.0) using Minimap2. The reads were then aligned against the diploid genome to understand the haplotype with the translocated chromosome X. After isolation of the reads to confirm the breakpoint sequence, the final marker chromosome X with the t(Xq;10q) was manually curated. The resulting fasta file was then validated by re-alignment of the reads that span the breakpoint for >99% identity (Supplementary Note 4).

### Annotation and analysis of centromere repetitive regions

Centromere estimation was done intersecting the outputs derived from RepeatMasker (https://github.com/rmhubley/RepeatMasker) ^51^ and HumAS-HMMER For AnVIL (https://github.com/fedorrik/HumAS-HMMER_for_AnVIL) to identify the position of the centromeres in the *de novo* RPE-1v1.0 assembly, and the HOR-monomer annotation was used to predict HOR Structural Variants (SVs) using StV script (https://github.com/fedorrik/stv). RepeatMasker was used with default parameters, and HumAS-HMMER as For AnVIL described in ^22^. To generate heatmaps showing the variation between centromeres, StainedGlass (v6.7.0) (https://github.com/mrvollger/StainedGlass) ^88^ was run with the following parameters: window=1000 and mm_f=10000. StVs specific for each haplotype were obtained using bedtools. HORs and StVs were visualized on IGV.

### OMICS data analysis

CUT&RUN reads from the publicly available dataset of hTERT RPE-1 cells (GSE132193) ^63^ were assessed for quality using FastQC (v0.11.9) (https://github.com/s-andrews/FastQC). CENP-A (SRA: SRR9201843) CUT&RUN reads and WGS Input (SRA: SRR9201844) of RPE-1 cells were aligned against multiple genomes: our *de novo* RPE-1 phased haplotypes (diploid mapping), the T2T-CHM13 whole-genome assembly v2.0 (haploid mapping), the HG38 (GRCh38.p14, haploid mapping) and the HG002v1.0 paternal and maternal haplotypes (diploid mapping). Bowtie2 (v2.4.4) (https://github.com/BenLangmead/bowtie2) ^89^ was used with the following parameters: bowtie2--end-to-end-x [index_reference] [read1.fastq] [read2.fastq] for paired-end data. The parameter end-to-end allows to search for alignments involving all the read characters instead of performing a local alignment. The resulting alignment files were filtered using SAMtools (v1.12) (https://github.com/samtools/samtools) ^90^ with FLAG score 2308 to avoid secondary and supplementary mapping of the reads, hence, preventing mapping biases in highly identical regions. Significantly enriched (removing FLAG 2308 and retaining peaks with q-value ≤0.00001) and high-confidence (removing FLAG 2308, considering MAPQ >20 and retaining peaks with q-value ≤0.00001) CENP-A peaks were determined using MACS3 (v3.0.0b1) (https://github.com/macs3-project/MACS), calculating the ratio between immunoprecipitated samples and background (WGS Input) with these parameters: -f BAMPE -B -g 3.03e9 -q 0.00001. High-confidence CENP-A positions were also identified using the mapping scores (MAPQ: 20, corresponding to the probability that the correct mapping to another location is 0.01) to identify reads that aligned uniquely to low-frequency repeat variants. The package karyoploteR (https://github.com/bernatgel/karyoploteR) ^91^ was used to generate the CENP-A density visualization across all chromosomes. Highly significant CENP-A peaks were computed using the internal function kpPlotDensity using a 500 bp window size and peaks called with a q-value ≤0.00001. For CENP-A high-confidence peaks, the window size was 50 bp and the q-value ≤0.00001.

### Methylation processing with Dorado

Using Dorado (v0.3.0) (https://github.com/nanoporetech/dorado), we downloaded the simplex basecalling model (dna_r10.4.1_e8.2_400bps_sup@v4.2.0) and we called the methylation profile with the default parameter --modified-bases 5mC and with --modified-bases-threshold 0.08. Dorado outputs directly the modified 5mC in the SAM/BAM files produced with mapping in parallel all the ONT reads against the diploid RPE-1v1.0 reference genome. It adds the MM (base modification methylation) and ML (base modification probability) tags that represent the relative position and probability that particular canonical bases have the requested modified bases. The output obtained with Dorado was then processed using Modkit (v0.2.0) (https://github.com/nanoporetech/modkit) with --filter-threshold 0.80 parameters to get a bedMethyl file with the annotation and the position of each single modification and its relative coverage. The DNA methylation profile was plotted using the internal function kpPlotDensity with a 5000 kb window size.

## DATA AVAILABILITY

Release of the RPE-1v1.0 reference genome has been approved for public access by Geron Corp. All scripts and resources generated and/or used in this project are publicly available at (https://github.com/GiuntaLab/RPE1).

## ACKNOWLEDGEMENTS

We thank Valentina Liguori and all members of the Giunta lab, as well as Alessandro Paiardini, Beniamino Trombetta, Fulvio Cruciani at University of Rome Sapienza, for helpful discussions and insights. We are especially indebt with Luigi Faino at Sapienza for help with HPC troubleshooting, key computational insights and advice. We are thankful to T2T and HPRC members for the kind assistance, especially Sergey Koren with Verkko, Nadolina Brajuka and members of Erich Jarvis lab for HiC data integration. We acknowledge Karen Miga, Glennis Logsdon and Jennifer Gerton for sharing information and useful discussions. We are grateful to Nicolas Altemose and Altemose lab members, as well as Riccardo Paone, Isabella Baldini, Juan Caceres and Noemi Di Sabatino at Dante Labs, and Margherita de Gaspari at Area Science Park, for generous sharing of insight, space and resources. We thank Alessandra Bonito Oliva at Rockefeller University, as well as Sai Swaroop Chittoor, Giulio Marano, Veronica Marselli and Alessia Spurio, for critical reading of the manuscript and providing feedback.

All computing was possible thanks to CINECA, and the Terastat2 HPC, with important support from Umberto Ferraro Petrillo and Edoardo Bompiani of the Statistics Department, University of Rome Sapienza.

This work was possible thanks to the Italian Foundation for Cancer Research (AIRC Start-Up Grant 2020 ID. # 25189) and the Rita Levi-Montalcini program from the Italian Ministry of University and Research (MIUR) to S.G., and the PON Ph.D. Fellowship to E.V. by the MIUR, Italy. G.F. was supported by NIH NHGRI Grant U01HG010971.

## AUTHOR CONTRIBUTIONS

S.G. conceived the idea underlying this work; S.G. and E.V. designed the study with key input from L.C.. E.V., and D.L. generated all sequencing data, with supervision from R.O. and M.C. for PacBio data. E.V. and L.C. performed all computational analyses. E.D.T., S.G. and F.P. designed and performed cytogenetic experiments, while E.V. performed all other experiments. S.G supervised experiments and analyses with key support from E.T. and G.F.. E.D.T. and S.G. designed and assembled all figures with help from L.C., E.V. and A.G.. S.G. wrote the manuscript together with E.T., L.C., E.V., E.D.T. with key editing from G.F. and A.G.. All authors have read and approved the manuscript.

## Supplemental information Supplemental Figure Legends

**Supplementary Figure 1.**
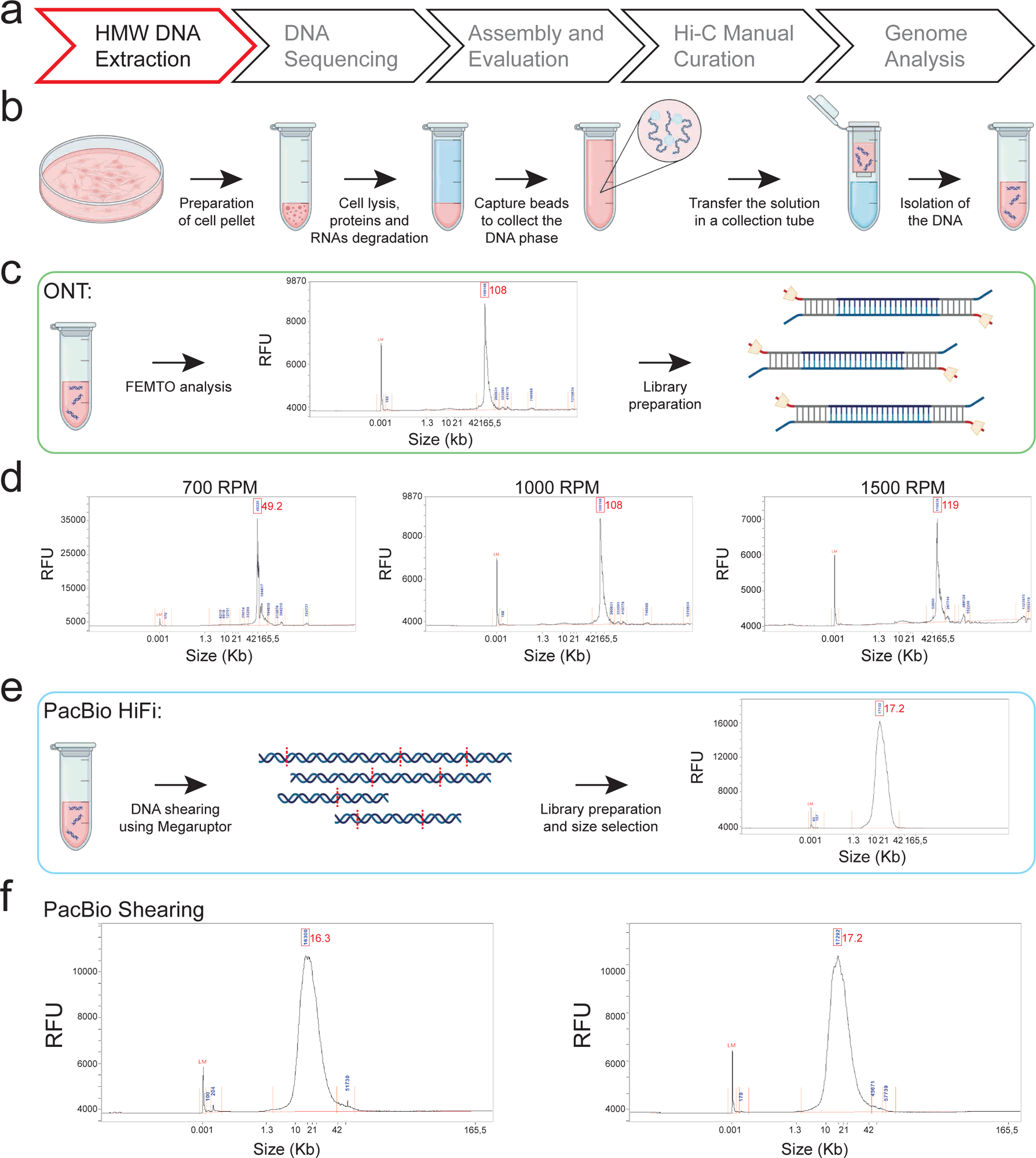
DNA extraction and libraries preparation. (a) Workflow applied to obtain de novo genome assembly, High and ultra-High Molecular Weight DNA extraction step (HMW and uHMW, respectively). (b) DNA was extracted from RPE-1 cell pellets according to a specific centrifugation speed, and beads were used to collect genomic DNA. (c) uHMW DNA enrichment was performed for ONT libraries. (d) Optimal length of extracted DNA was evaluated using FEMTO pulse. (e) HMW DNA extraction was performed for PacBio libraries. (f) The final library length for PacBio was evaluated using FEMTO pulse.

**Supplementary Figure 2.**
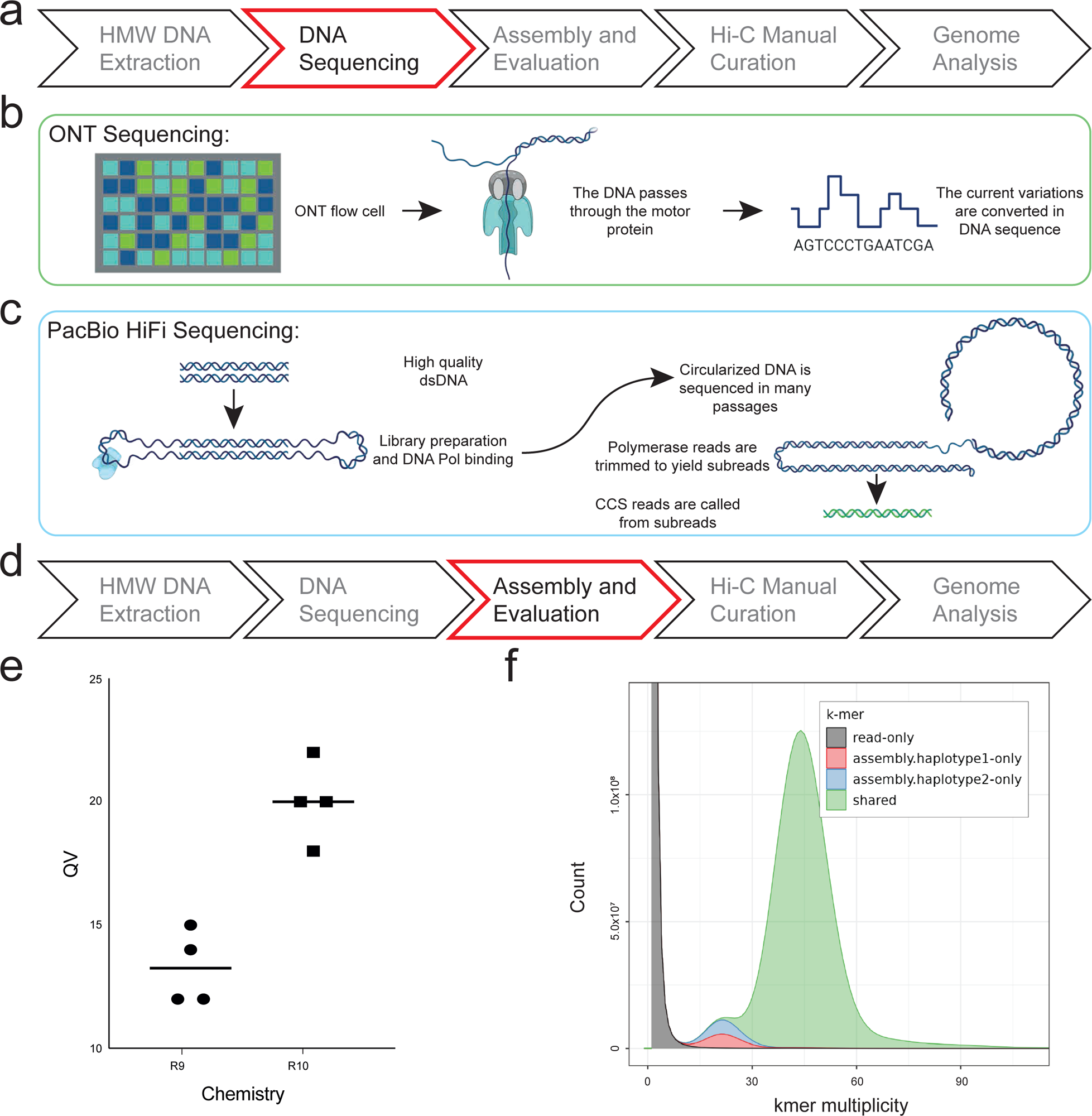
Long and ultra-long read sequencing. (a) Workflow applied to obtain de novo diploid genome assembly, DNA sequencing step. (b) ONT sequencing was performed using PromethION and R10.4 flow cells. (c) PacBio sequencing was performed using Sequel IIe and 8 million Zero Mode Waves flow cells (ZMWs), producing Circular Consensus High Fidelity reads (CCS, HiFi reads). (d) Assembly and quality evaluation step. (e) R10.4 chemistry increased ONT base calling accuracy from 98 to 99% compared to R9. (f) K-mer spectra analysis performed with *Merqury* highlights the quality of the final assembly by showing two separate peaks for both haplotypes and one shared peak due to homozygous regions.

**Supplementary Figure 3.**
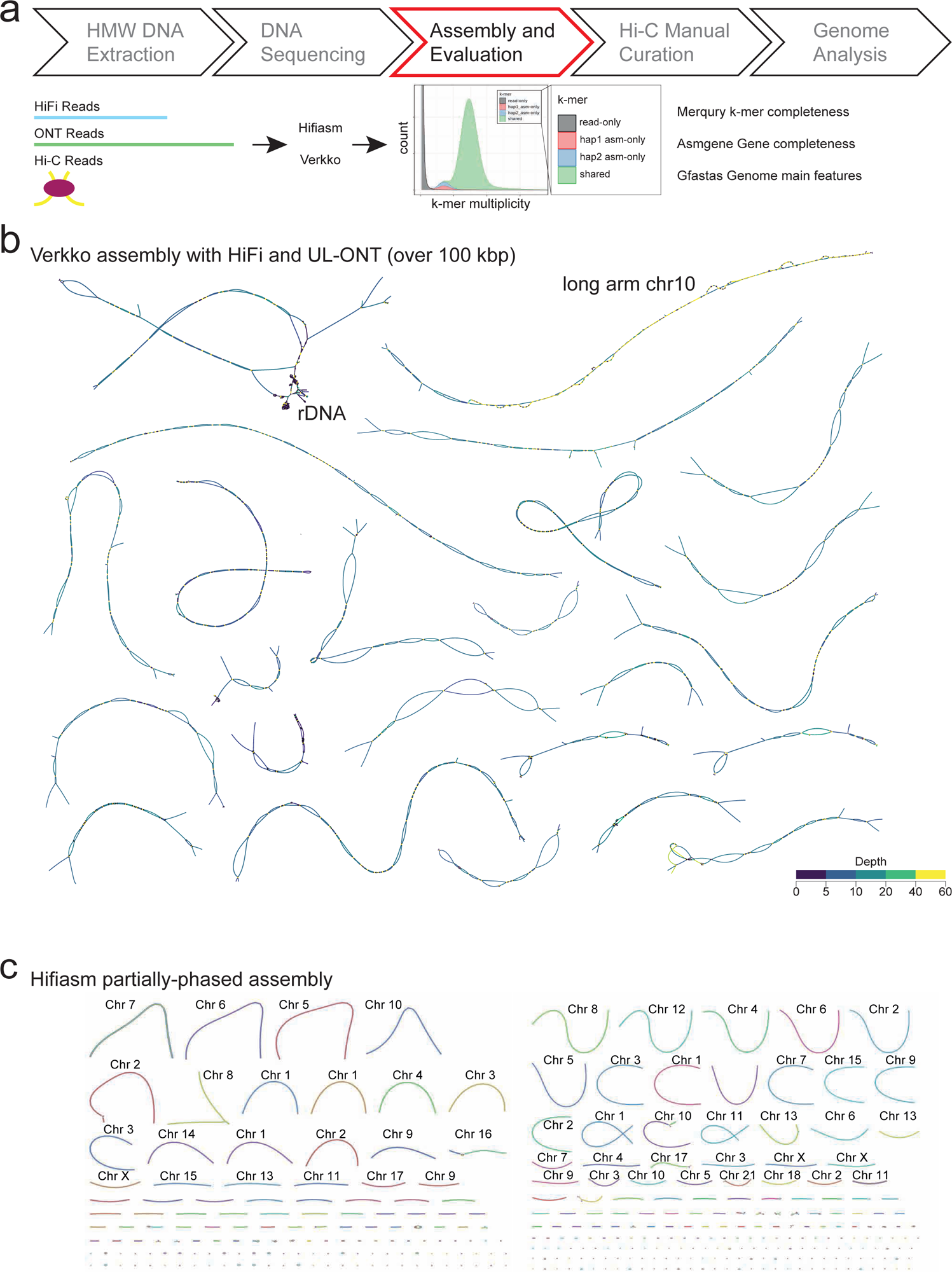
*De novo* diploid assembly. (a) Workflow applied to obtain *de novo* diploid genome assembly, assembly and evaluation steps. (b) The first assembly was generated using PacBio HiFi and ultra-long (UL) ONT reads (> 100 kb). The string graph obtained using Verkko tool was visualized with Bandage. Graph nodes are colored by read depth, with the yellow that represents the highest value. The extra copy of the duplicated q-arm of chromosome 10 is completely colored in yellow confirming the duplicated state of this region. (c) Hifiasm was used with only HiFi reads, resulting in a string graph with partially-phased haplotypes visualized with Bandage.

**Supplementary Figure 4.**
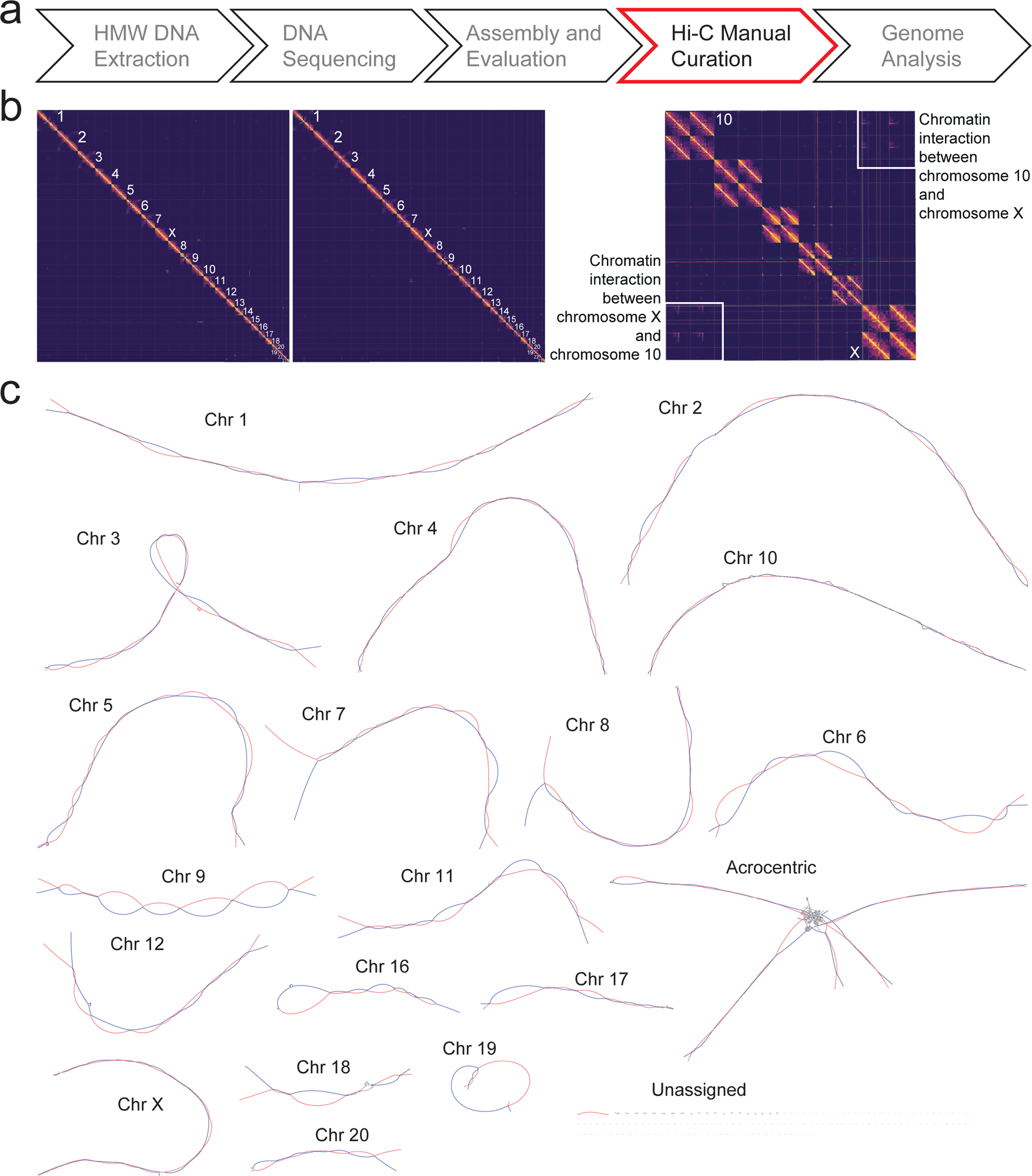
Phasing of the *de novo* diploid assembly. (a) Workflow applied to obtain *de novo* diploid genome assembly with Hi-C data integration and curation. (b) Hi-C 3D contact maps, visualized using Pretext, show the two haplotypes at chromosome-level resolution. Hi-C data confirm chromatin interaction between chromosome 10 and chromosome X (right). (c) The string graph of the final phased assembly was visualized using Bandage. The haplotypes are colored in red and blue. Short arms of chr13, chr14, chr15, chr21, chr22 are tangled in correspondence of the rDNA arrays.

**Supplementary Figure 5.**
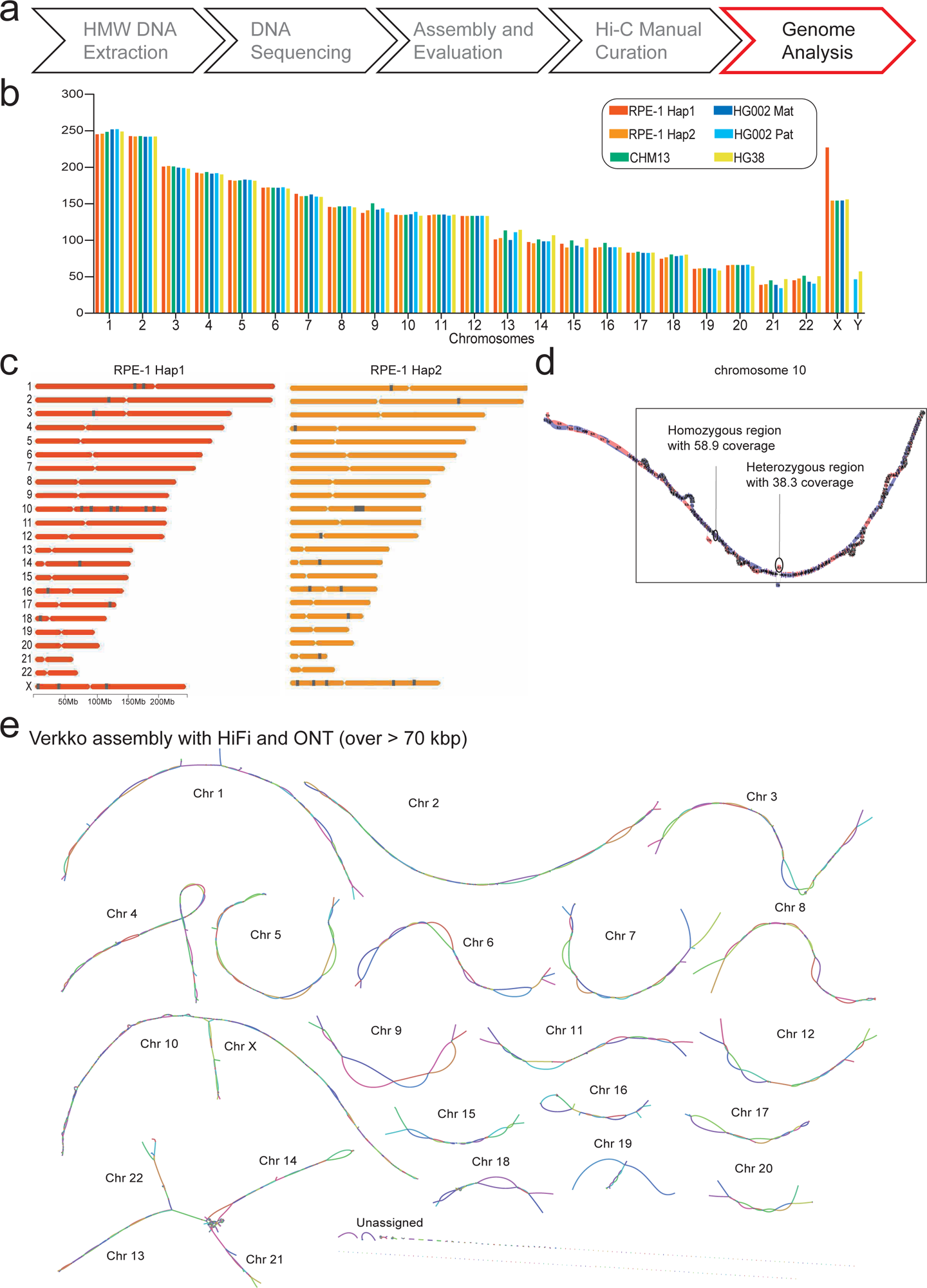
Assembly quality evaluation for final diploid phased genome. (a) Workflow applied to obtain de novo diploid genome assembly, genome analysis steps. (b) Chromosome lengths of RPE-1 are comparable with the previously assembled genomes, excluding chromosome X of Hap 1 which harbors the structural variant characteristic of the RPE-1 cell line, that is a translocation of part of chromosome 10. (c) The presence of the duplicated long arm region of chromosome 10 on chromosome X led to an increased number of gaps compared to the other chromosomes. (d) The extra copy of chromosome 10 is also shown in the string graph as an increase in coverage in homozygous and heterozygous regions. (e) Verkko assembly using all ONT reads resulted in the final graph with fused chromosomes 10 and X. The graph is visualized with Bandage.

**Supplementary Figure 6.**
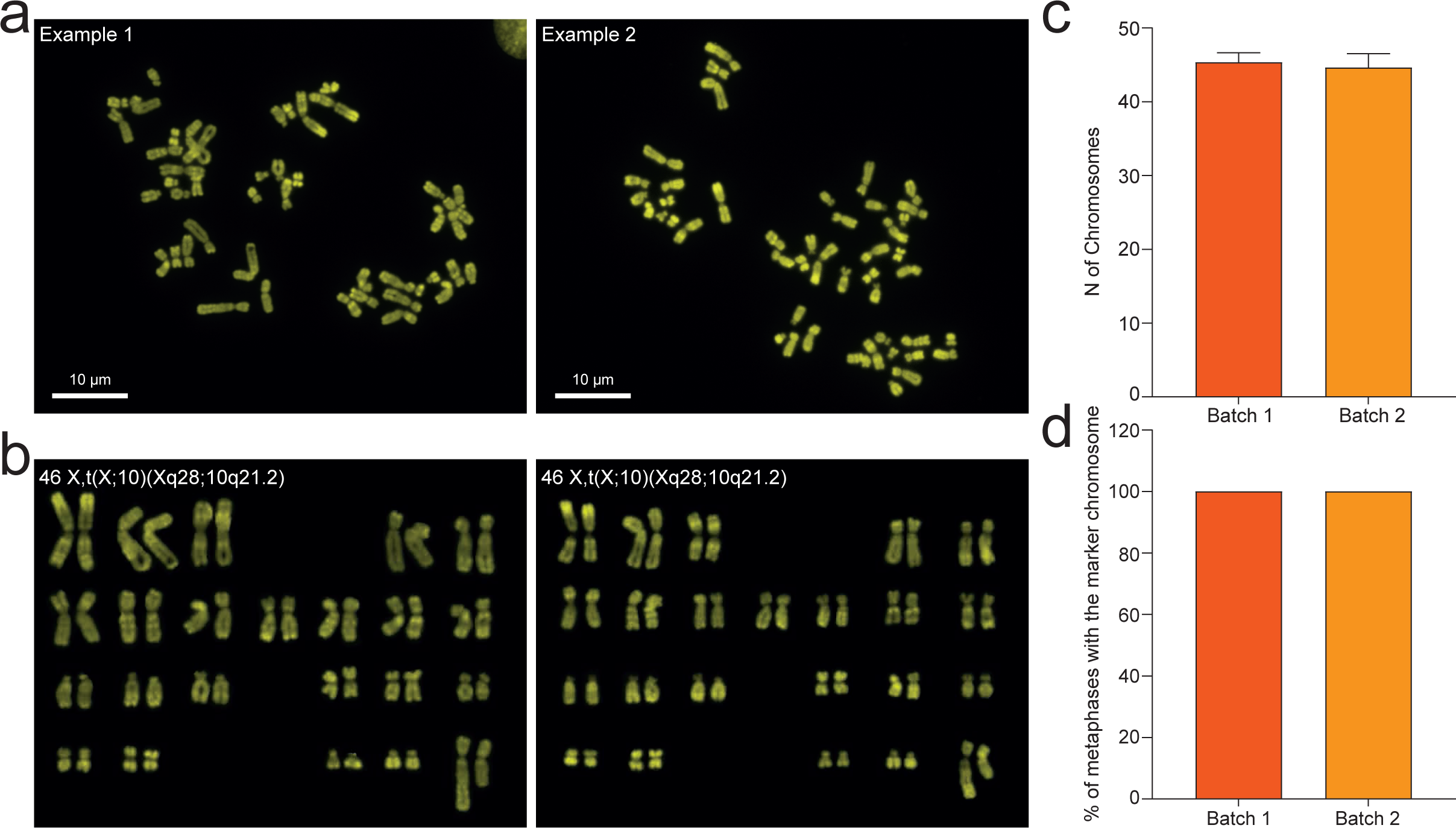
Karyotype of the RPE-1 cell line. a) Chromomycin A3 staining was used to obtain an R-banding of the chromosomes of the RPE-1 cell line. The scale bar is 10 μm. b) The banding pattern allowed the identification of the chromosomes, that were then ordered to obtain a karyogram. c) Metaphase analysis revealed that RPE-1 cells are in a diploid state. This result was confirmed in two different batches of cells obtained from ATCC or another laboratory. d) The presence of the marker chromosome of the RPE-1 cell line, t(X;10) (Xq28;10q21.2), was confirmed in all metaphases analyzed.

**Supplementary Figure 7.**
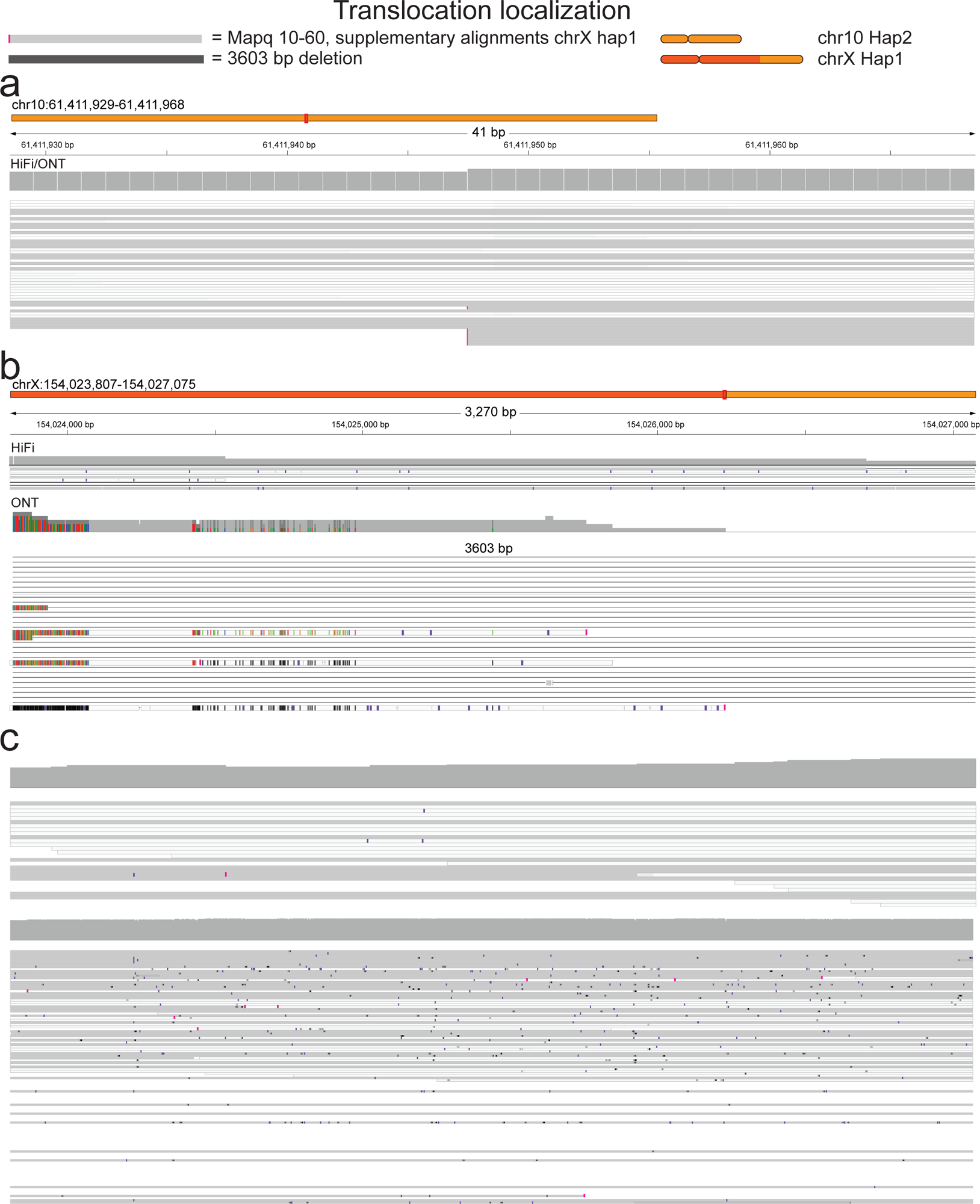
Marker chromosome breakpoint resolution and manual curation. (a) IGV visualization of the HiFi and ONT read alignments against the RPE-1 diploid genome. The alignment of chromosome 10 for Hap 2 highlighted an interruption in read coverage in position 61640000 bp, showing reads with 10-60 mapping quality and supplementary alignment at chrX:154027075. Notably, these supplementary alignments belong only to chromosome X of Hap 1. (b) HiFi and ONT reads were aligned against chromosome X harboring the additional sequence coming from the long arm of chromosome 10. The alignment shows a telomere region with a 3603 bp deletion at the telomeric region. (c) HiFi and ONT reads alignment against the sequence of chromosome X deleted of these 3603 bp in the telomeric region showed a complete alignment through the break point.

**Supplementary Figure 8.**
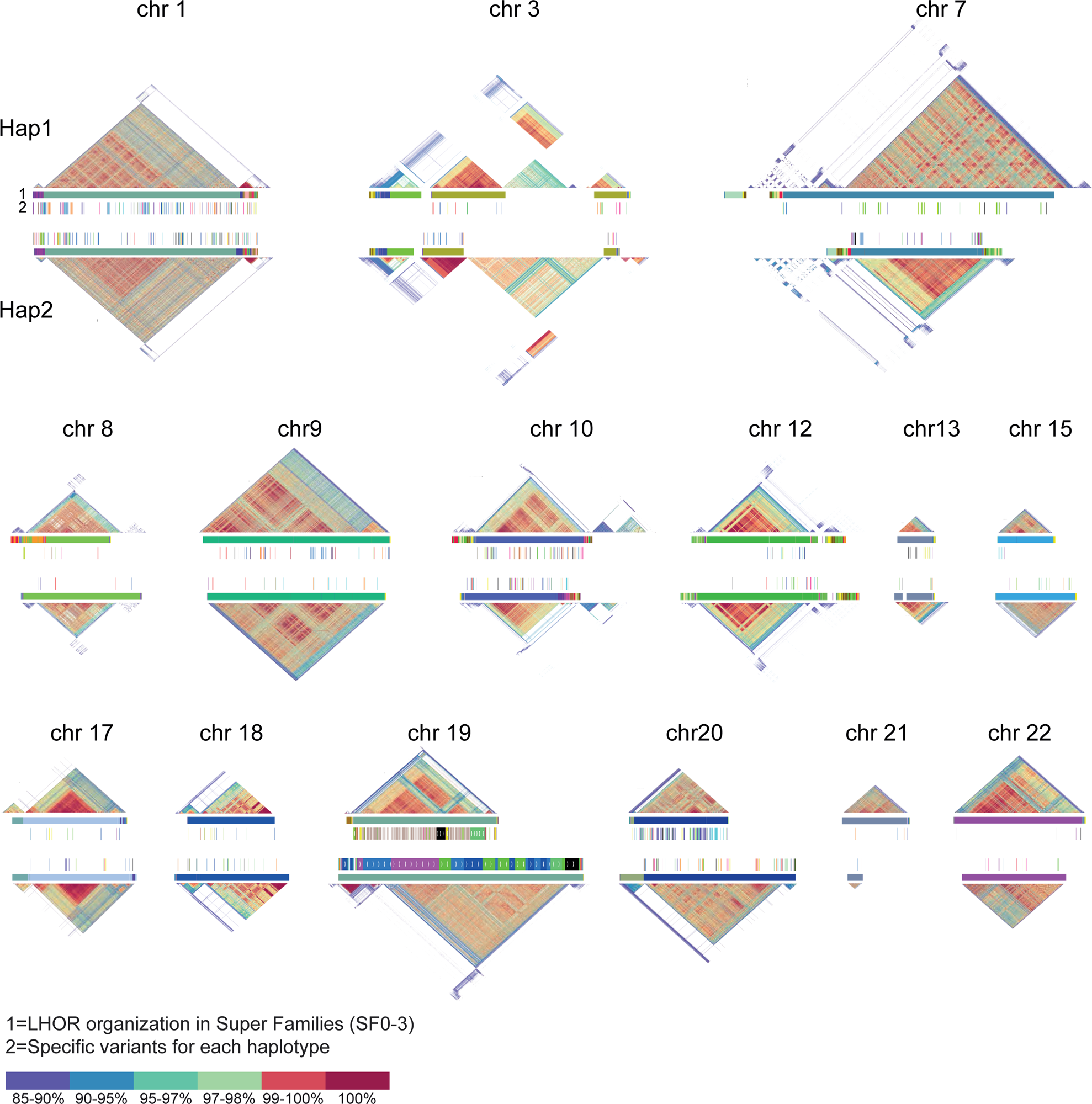
Centromere variation. Pairwise similarity heatmaps, created using StainedGlass, of centromeres and LHORs (Live High Order Repeats) for all chromosomes. Chromosomes 7 and 21 show differences in centromere size, while centromeres of chromosomes 9 and 19 show different HORs organization. HOR structural variants (SVs), which belong specifically to Hap 1 or 2, are also shown.

**Supplementary Figure 9.**
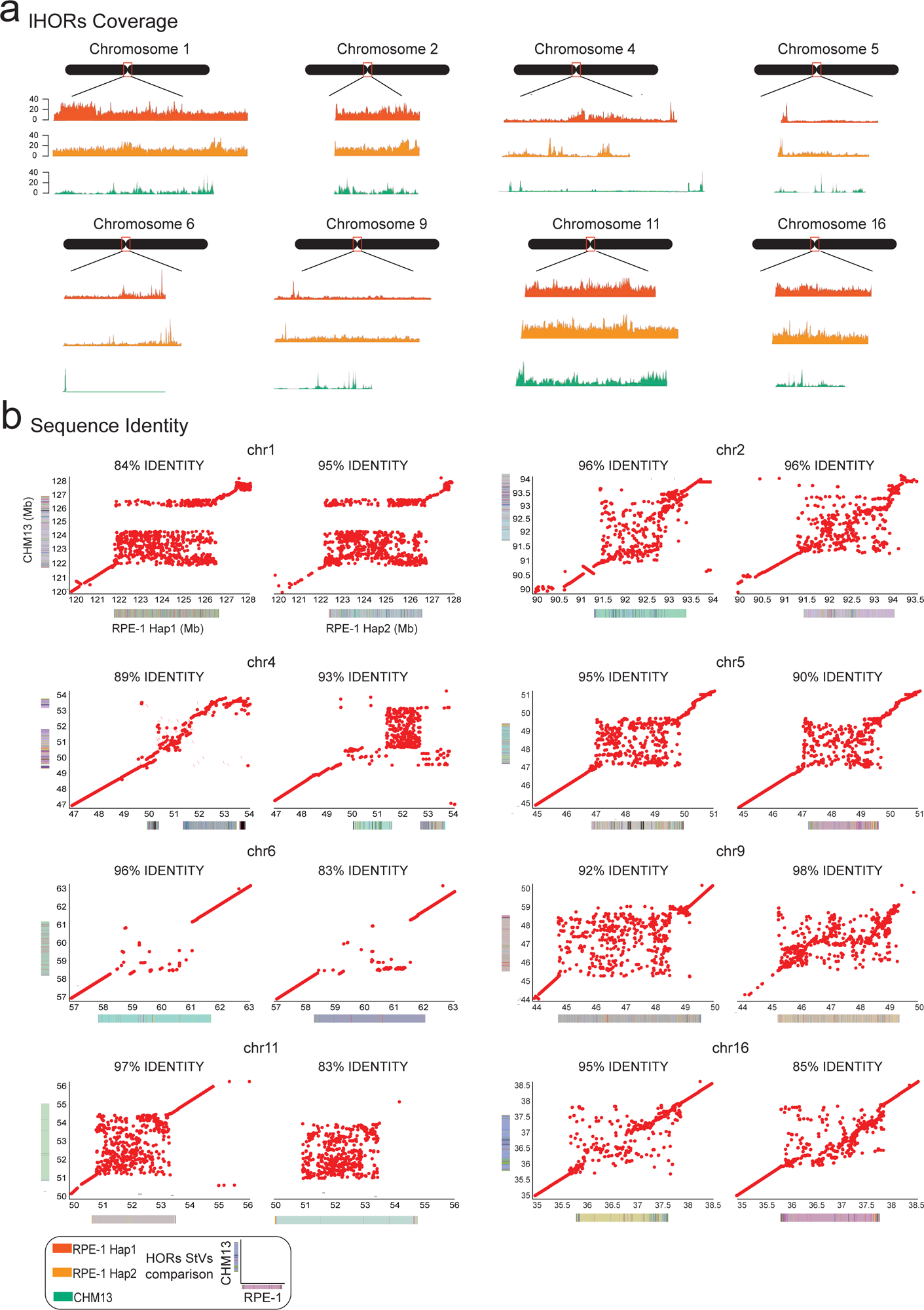
Zooming in on centromere regions of HiFi RPE-1 reads alignment against RPE-1 Hap 1, 2, and CHM13. (a) RPE-1 reads aligned against CHM13 show a complete absence of matched reads for chromosome 4 and chromosome 6, suggesting completely different LHORs structures and organizations. (b) Sequence identity, computed with MashMap, was evaluated between each haplotype and CHM13 genome. The lowest values were observed in the comparison between chromosome 1 CHM13 versus RPE-1 Hap 1, and chromosome 6 and chromosome 11 CHM13 against RPE-1 Hap 2.

**Supplementary Figure 10.**
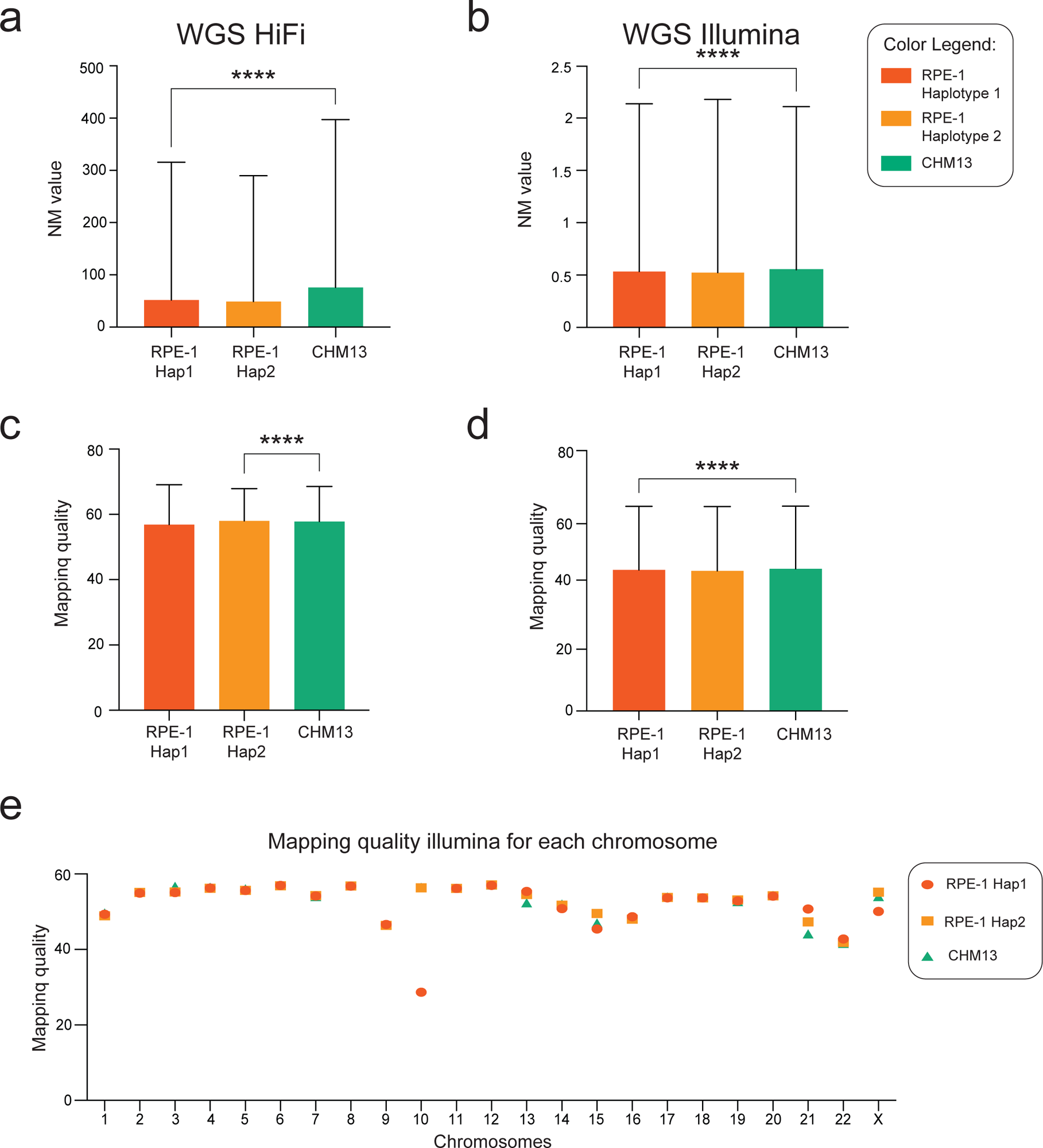
The mapping quality and NM values for HiFi and Illumina reads across the whole genome. (a-b) HiFi and Illumina read alignment exhibit a statistically significant decrease in NM values (edit distance) when RPE-1 Hap 1 and 2 are used as a reference. (c-d) Mapping quality showed a significant increase for Hap 2 compared to CHM13, and a decrease comparing Hap 1 and CHM13. (e) The analysis of the mapping quality for each chromosome explains its decrease when Hap 1 is compared to CHM13. The duplication in RPE-1 Hap 1 leads to a decrease in mapping quality due to the presence of two copies of the same sequence for chromosome 10.

**Supplementary Figure 11.**
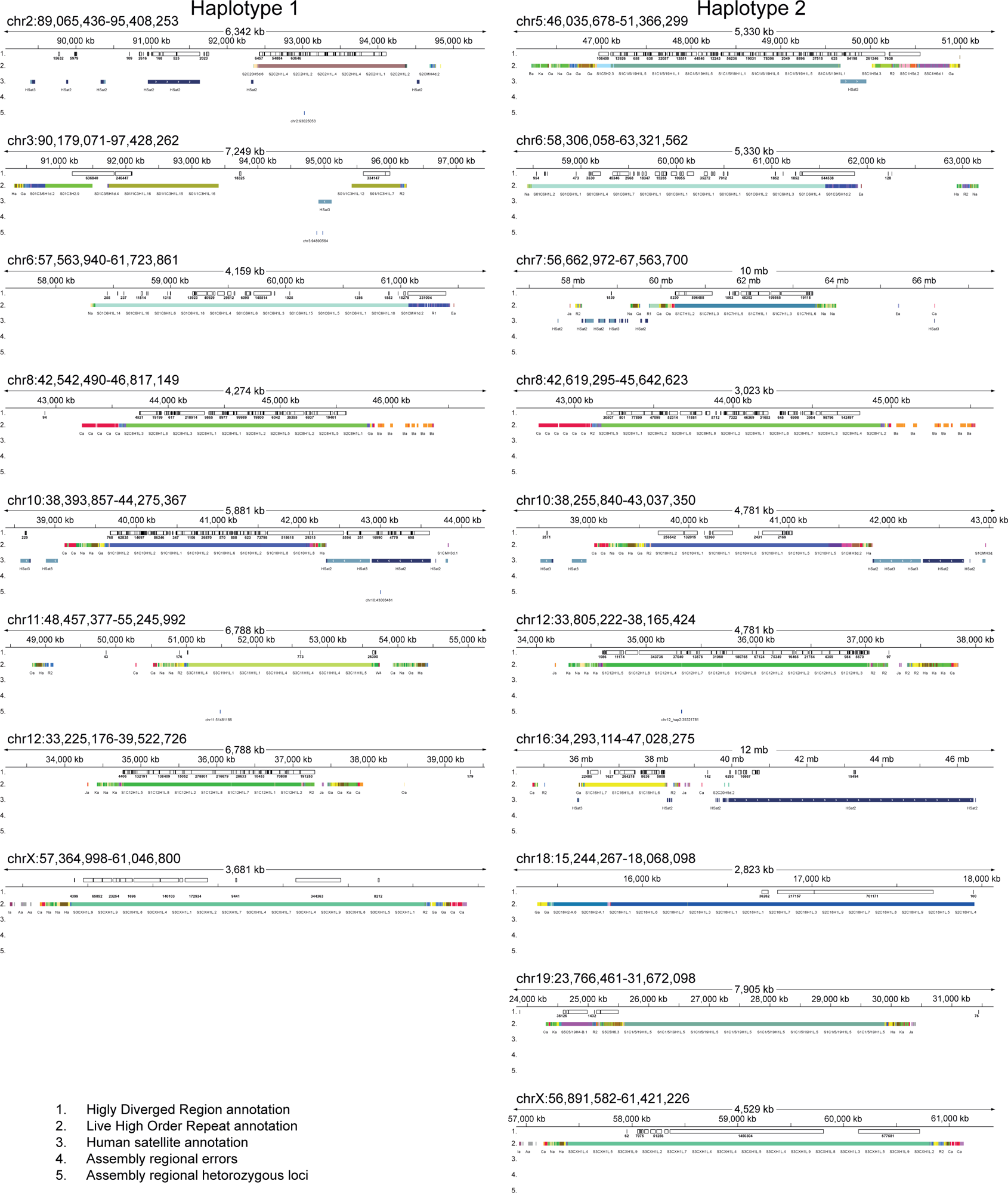
The isogenomic reference genome improves reads alignment. RPE-1 Hap 1 and Hap 2 were compared with CHM13. Analyzing the SVs obtained with SyRI, a total of 37% and 57% of HDRs belong to centromeric regions when Hap 1 and Hap 2 were compared to CHM13. Chromosomes with 50% or more of HDRs belonging to centromeric regions were analyzed to exclude that the high divergence were due to assembly errors. HDRs of chromosomes 3, 8, and 12 for Hap 1, and chromosomes 10, 12, and 18 for Hap 2 are located almost exclusively in centromere regions for a total of 90% of their length. *Craq* tool was used to evaluate the total number of errors and heterozygous bases in the assembly. The output shows complete absence of regional assembly errors, but only the presence of structural heterozygous variants. The final IGV visualization indicates HDRs, HORs, Hsat, errors, and heterozygous positions for the RPE1v1.0 reference genome assembly.

## References

1. Nurk, S. et al. The complete sequence of a human genome. Science 376, 44–53 (2022).

2. Jarvis, E. D. et al. Semi-automated assembly of high-quality diploid human reference genomes. Nature 611, 519–531 (2022).

3. Liao, W.-W. et al. A draft human pangenome reference. Nature 617, 312–324 (2023).

4. Chao, K.-H., Zimin, A. V., Pertea, M. & Salzberg, S. L. The first gapless, reference-quality, fully annotated genome from a Southern Han Chinese individual. G3 Bethesda Md 13, jkac321 (2023).

5. Yang, C. et al. The complete and fully-phased diploid genome of a male Han Chinese. Cell Res. 33, 745–761 (2023).

6. He, Y. et al. T2T-YAO: A Telomere-to-telomere Assembled Diploid Reference Genome for Han Chinese. Genomics Proteomics Bioinformatics S1672–0229(23)00100–6 (2023).

7. Wang, T. et al. The Human Pangenome Project: a global resource to map genomic diversity. Nature 604, 437–446 (2022).

8. Yang, X. et al. Characterization of large-scale genomic differences in the first complete human genome. Genome Biol. 24, 157 (2023).

9. Karczewski, K. J. et al. The mutational constraint spectrum quantified from variation in 141,456 humans. Nature 581, 434–443 (2020).

10. Jónsson, H. et al. Whole genome characterization of sequence diversity of 15,220 Icelanders. Sci. Data 4, 170115 (2017).

11. Seplyarskiy, V. B. et al. Population sequencing data reveal a compendium of mutational processes in the human germ line. Science 373, 1030–1035 (2021).

12. Cheng, H., Concepcion, G. T., Feng, X., Zhang, H. & Li, H. Haplotype-resolved de novo assembly using phased assembly graphs with hifiasm. Nat. Methods 18, 170–175 (2021).

13. The International HapMap Consortium. A haplotype map of the human genome. Nature 437, 1299–1320 (2005).

14. Chiang, C. et al. The impact of structural variation on human gene expression. Nat. Genet. 49, 692–699 (2017).

15. Pollen, A. A., Kilik, U., Lowe, C. B. & Camp, J. G. Human-specific genetics: new tools to explore the molecular and cellular basis of human evolution. Nat. Rev. Genet. 24, 687–711 (2023).

16. Eichler, E. E. Genetic Variation, Comparative Genomics, and the Diagnosis of Disease. N. Engl. J. Med. 381, 64–74 (2019).

17. Cancellieri, S. et al. Human genetic diversity alters off-target outcomes of therapeutic gene editing. Nat. Genet. 55, 34–43 (2023).

18. Garrison, E. et al. Variation graph toolkit improves read mapping by representing genetic variation in the reference. Nat. Biotechnol. 36, 875–879 (2018).

19. Hayden, K. E. Human centromere genomics: now it’s personal. Chromosome Res. 20, 621– 633 (2012).

20. Schueler, M. G. & Sullivan, B. A. Structural and functional dynamics of human centromeric chromatin. Annu. Rev. Genomics Hum. Genet. 7, 301–313 (2006).

21. Willard, H. F. Chromosome-specific organization of human alpha satellite DNA. Am. J. Hum. Genet. 37, 524–532 (1985).

22. Altemose, N. et al. Complete genomic and epigenetic maps of human centromeres. Science 376, eabl4178 (2022).

23. McKinley, K. L. & Cheeseman, I. M. The molecular basis for centromere identity and function. Nat. Rev. Mol. Cell Biol. 17, 16–29 (2016).

24. Sullivan, K. F. A solid foundation: functional specialization of centromeric chromatin. Curr. Opin. Genet. Dev. 11, 182–188 (2001).

25. Gershman, A. et al. Epigenetic patterns in a complete human genome. Science 376, eabj5089 (2022).

26. Chaisson, M. J. P. et al. Resolving the complexity of the human genome using single-molecule sequencing. Nature 517, 608–611 (2015).

27. Nurk, S. et al. HiCanu: accurate assembly of segmental duplications, satellites, and allelic variants from high-fidelity long reads. Genome Res. gr.263566.120 (2020) doi:10.1101/gr.263566.120.

28. Vollger, M. R. et al. Segmental duplications and their variation in a complete human genome. Science 376, eabj6965 (2022).

29. Logsdon, G. A. & Eichler, E. E. The Dynamic Structure and Rapid Evolution of Human Centromeric Satellite DNA. Genes 14, 92 (2022).

30. Porubsky, D. et al. Inversion polymorphism in a complete human genome assembly. Genome Biol. 24, 100 (2023).

31. Logsdon, G. A., et al. The variation and evolution of complete human centromeres. http://biorxiv.org/lookup/doi/10.1101/2023.05.30.542849 (2023) doi:10.1101/2023.05.30.542849.

32. Steinberg, K. M. et al. Single haplotype assembly of the human genome from a hydatidiform mole. Genome Res. 24, 2066–2076 (2014).

33. Vollger, M. R. et al. Long-read sequence and assembly of segmental duplications. Nat. Methods 16, 88–94 (2019).

34. Logsdon, G. A. et al. The structure, function and evolution of a complete human chromosome 8. Nature 593, 101–107 (2021).

35. https://www.atcc.org/products/crl-4000. hTERT RPE-1. https://www.atcc.org/products/crl-4000.

36. Cheng, H. et al. Haplotype-resolved assembly of diploid genomes without parental data. Nat. Biotechnol. 40, 1332–1335 (2022).

37. Rautiainen, M. et al. Telomere-to-telomere assembly of diploid chromosomes with Verkko. Nat. Biotechnol. 41, 1474–1482 (2023).

38. Rhie, A., Walenz, B. P., Koren, S. & Phillippy, A. M. Merqury: reference-free quality, completeness, and phasing assessment for genome assemblies. Genome Biol. 21, 245 (2020).

39. Simão, F. A., Waterhouse, R. M., Ioannidis, P., Kriventseva, E. V. & Zdobnov, E. M. BUSCO: assessing genome assembly and annotation completeness with single-copy orthologs. Bioinformatics 31, 3210–3212 (2015).

40. Li, H. Minimap2: pairwise alignment for nucleotide sequences. Bioinformatics 34, 3094– 3100 (2018).

41. Li, K., Xu, P., Wang, J., Yi, X. & Jiao, Y. Identification of errors in draft genome assemblies at single-nucleotide resolution for quality assessment and improvement. Nat. Commun. 14, 6556 (2023).

42. Giunta, S. et al. CENP-A chromatin prevents replication stress at centromeres to avoid structural aneuploidy. Proc. Natl. Acad. Sci. U. S. A. 118, e2015634118 (2021).

43. Giunta, S. & Funabiki, H. Integrity of the human centromere DNA repeats is protected by CENP-A, CENP-C, and CENP-T. Proc. Natl. Acad. Sci. U. S. A. 114, 1928–1933 (2017).

44. Janssen, A., van der Burg, M., Szuhai, K., Kops, G. J. P. L. & Medema, R. H. Chromosome segregation errors as a cause of DNA damage and structural chromosome aberrations. Science 333, 1895–1898 (2011).

45. Soto, M. et al. p53 Prohibits Propagation of Chromosome Segregation Errors that Produce Structural Aneuploidies. Cell Rep. 19, 2423–2431 (2017).

46. Tourdot, R. W., Brunette, G. J., Pinto, R. A. & Zhang, C.-Z. Determination of complete chromosomal haplotypes by bulk DNA sequencing. Genome Biol. 22, 139 (2021).

47. Choo, K. H. Why is the centromere so cold? Genome Res. 8, 81–82 (1998).

48. Balzano, E. & Giunta, S. Centromeres under Pressure: Evolutionary Innovation in Conflict with Conserved Function. Genes 11, 912 (2020).

49. Goel, M., Sun, H., Jiao, W.-B. & Schneeberger, K. SyRI: finding genomic rearrangements and local sequence differences from whole-genome assemblies. Genome Biol. 20, 277 (2019).

50. Trowsdale, J. & Knight, J. C. Major histocompatibility complex genomics and human disease. Annu. Rev. Genomics Hum. Genet. 14, 301–323 (2013).

51. Smit, Hubley and Green. https://repeatmasker.org. (2013).

52. Maloney, K. A. et al. Functional epialleles at an endogenous human centromere. Proc. Natl. Acad. Sci. 109, 13704–13709 (2012).

53. Antonarakis, S. E. Parental origin of the extra chromosome in trisomy 21 as indicated by analysis of DNA polymorphisms. Down Syndrome Collaborative Group. N. Engl. J. Med. 324, 872– 876 (1991).

54. Levy-Sakin, M. et al. Genome maps across 26 human populations reveal population-specific patterns of structural variation. Nat. Commun. 10, 1025 (2019).

55. Delcher, A. L. et al. Alignment of whole genomes. Nucleic Acids Res. 27, 2369–2376 (1999).

56. Paten, B. et al. Cactus: Algorithms for genome multiple sequence alignment. Genome Res. 21, 1512–1528 (2011).

57. Yuan, S. & Qin, Z. Read-mapping using personalized diploid reference genome for RNA sequencing data reduced bias for detecting allele-specific expression. IEEE Int. Conf. Bioinforma. Biomed. Workshop IEEE Int. Conf. Bioinforma. Biomed. 2012, 718–724 (2012).

58. Paten, B., Novak, A. M., Eizenga, J. M. & Garrison, E. Genome graphs and the evolution of genome inference. Genome Res. 27, 665–676 (2017).

59. Altemose, N. et al. DiMeLo-seq: a long-read, single-molecule method for mapping protein-DNA interactions genome wide. Nat. Methods 19, 711–723 (2022).

60. Nechemia-Arbely, Y. et al. DNA replication acts as an error correction mechanism to maintain centromere identity by restricting CENP-A to centromeres. Nat. Cell Biol. 21, 743–754 (2019).

61. Saayman, X., Graham, E., Nathan, W. J., Nussenzweig, A. & Esashi, F. Centromeres as universal hotspots of DNA breakage, driving RAD51-mediated recombination during quiescence. Mol. Cell 83, 523–538.e7 (2023).

62. Mahlke, M. A., Lumerman, L., Ly, P. & Nechemia-Arbely, Y. Epigenetic centromere identity is precisely maintained through DNA replication but is uniquely specified among human cells. Life Sci. Alliance 6, e202201807 (2023).

63. Dumont, M. et al. Human chromosome-specific aneuploidy is influenced by DNA-dependent centromeric features. EMBO J. 39, e102924 (2020).

64. Rhie, A. et al. The complete sequence of a human Y chromosome. Nature 621, 344–354 (2023).

65. Nechemia-Arbely, Y. et al. Human centromeric CENP-A chromatin is a homotypic, octameric nucleosome at all cell cycle points. J. Cell Biol. 216, 607–621 (2017).

66. Gamba, R. et al. Enrichment of centromeric DNA from human cells. PLoS Genet. 18, e1010306 (2022).

67. Cappelletti, E. et al. The localization of centromere protein A is conserved among tissues. *Commun*. Biol. 6, 963 (2023).

68. Fachinetti, D. et al. A two-step mechanism for epigenetic specification of centromere identity and function. Nat. Cell Biol. 15, 1056–1066 (2013).

69. Thakur, J. & Henikoff, S. CENPT bridges adjacent CENPA nucleosomes on young human α-satellite dimers. Genome Res. 26, 1178–1187 (2016).

70. Bosco, N. et al. KaryoCreate: A CRISPR-based technology to study chromosome-specific aneuploidy by targeting human centromeres. Cell 186, 1985–2001.e19 (2023).

71. Schneider, V. A. et al. Evaluation of GRCh38 and de novo haploid genome assemblies demonstrates the enduring quality of the reference assembly. Genome Res. 27, 849–864 (2017).

72. Li, R. et al. Building the sequence map of the human pan-genome. Nat. Biotechnol. 28, 57– 63 (2010).

73. Chen, S., Zhou, Y., Chen, Y. & Gu, J. fastp: an ultra-fast all-in-one FASTQ preprocessor. Bioinformatics 34, i884–i890 (2018).

74. Andrews S. FastQC A Quality Control tool for High Throughput Sequence Data. Available online at: http://www.bioinformatics.babraham.ac.uk/projects/fastqc. https://www.bioinformatics.babraham.ac.uk/projects/fastqc/ (2010).

75. Martin, M. Cutadapt removes adapter sequences from high-throughput sequencing reads. EMBnet.journal 17, 10–12 (2011).

76. Ranallo-Benavidez, T. R., Jaron, K. S. & Schatz, M. C. GenomeScope 2.0 and Smudgeplot for reference-free profiling of polyploid genomes. Nat. Commun. 11, 1432 (2020).

77. Larivière, D. et al. Scalable, accessible, and reproducible reference genome assembly and evaluation in Galaxy. 2023.06.28.546576 Preprint at 10.1101/2023.06.28.546576 (2023).

78. Jain, C., Koren, S., Dilthey, A., Phillippy, A. M. & Aluru, S. A fast adaptive algorithm for computing whole-genome homology maps. Bioinformatics 34, i748–i756 (2018).

79. Marçais, G. et al. MUMmer4: A fast and versatile genome alignment system. PLOS Comput. Biol. 14, e1005944 (2018).

80. Lin, H.-N. & Hsu, W.-L. GSAlign: an efficient sequence alignment tool for intra-species genomes. BMC Genomics 21, 182 (2020).

81. Wick, R. R., Schultz, M. B., Zobel, J. & Holt, K. E. Bandage: interactive visualization of de novo genome assemblies. Bioinformatics 31, 3350–3352 (2015).

82. Limasset, A., Cazaux, B., Rivals, E. & Peterlongo, P. Read mapping on de Bruijn graphs. BMC Bioinformatics 17, 237 (2016).

83. Formenti, G. et al. Gfastats: conversion, evaluation and manipulation of genome sequences using assembly graphs. Bioinformatics 38, 4214–4216 (2022).

84. Gurevich, A., Saveliev, V., Vyahhi, N. & Tesler, G. QUAST: quality assessment tool for genome assemblies. Bioinformatics 29, 1072–1075 (2013).

85. Thorvaldsdóttir, H., Jt, R. & Jp, M. Integrative Genomics Viewer (IGV): high-performance genomics data visualization and exploration. Brief. Bioinform. 14, (2013).

86. Li, H. New strategies to improve minimap2 alignment accuracy. Bioinformatics 37, 4572– 4574 (2021).

87. Goel, M. & Schneeberger, K. plotsr: visualizing structural similarities and rearrangements between multiple genomes. Bioinformatics 38, 2922–2926 (2022).

88. Vollger, M. R., Kerpedjiev, P., Phillippy, A. M. & Eichler, E. E. StainedGlass: interactive visualization of massive tandem repeat structures with identity heatmaps. Bioinforma. Oxf. Engl. 38, 2049–2051 (2022).

89. Langmead, B. & Salzberg, S. L. Fast gapped-read alignment with Bowtie 2. Nat. Methods 9, 357–359 (2012).

90. Li, H. et al. The Sequence Alignment/Map format and SAMtools. Bioinformatics 25, 2078– 2079 (2009).

91. Gel, B. & Serra, E. karyoploteR: an R/Bioconductor package to plot customizable genomes displaying arbitrary data. Bioinforma. Oxf. Engl. 33, 3088–3090 (2017).

